# A discovery and replication study of dyslexia does not reveal reproducible gray matter volume differences

**DOI:** 10.64898/2026.05.05.722925

**Authors:** Alison K. Schug, Iria S. Gutierrez-Schieferl, Guinevere F. Eden

## Abstract

Two decades of research have provided evidence for gray matter volume (GMV) differences in developmental dyslexia (or reading disability, RD) in the left perisylvian cortex. However, there are concerns about result inconsistencies, likely attributable to small sample sizes, lenient statistical thresholds, and insufficient accounting for demographic variables and global GMV (Ramus et al., 2018). To address these concerns, we conducted a Discovery and Replication Study (N=262) using data from the Adolescent Brain Cognitive Development Study. We found GMV differences between the RD and Control Groups did not replicate across the Discovery and Replication Studies using voxel-based morphometry (VBM) in Statistical Parametric Mapping (SPM), and that a more conservative threshold yielded far fewer results. We then conducted Reproducibility Studies and first found that when using surface-based morphometry in FreeSurfer instead of VBM, the Discovery and the Replication Study results again failed to converge. Second, we combined all groups in a factorial VBM/SPM analysis and the interaction analysis provided quantitative confirmation for diverging between-group difference results across the two studies. Third, we tested for the role of covariates of no interest and found that when total GMV is not controlled for, this divergence dissipates and group differences in RD (main effect of Reading Ability) are amplified. In conclusion, replication of GMV differences in RD is low, even when using large, well-matched groups, and analyses approaches play a modulating role. As such, results from prior studies using lenient statistical thresholds and not accounting for total GMV should therefore be viewed with caution.

## 1 Introduction

Since the earliest studies into developmental dyslexia (also referred to as reading disability, RD, and used here interchangeably), neuroanatomical aberrations have been a central focus (Drake, 1968). Over the last two decades there have been numerous reports of less gray matter volume (GMV) in dyslexia, many of which focus on left-hemisphere perisylvian cortical regions known to be involved in language. Meta-analyses are a helpful tool for revealing convergence across study results and, when applied to identify the most salient GMV differences in dyslexia relative to controls (in users of the alphabet), found less GMV in bilateral temporoparietal, left occipitotemporal, and prefrontal cortices as well as the cerebellum (Eckert et al., 2016; Linkersdörfer et al., 2012; Richlan et al., 2012). However, as pointed out by Ramus and colleagues (2018), the consistency among the results is not all that strong. For example, while these three meta-analyses overall represent a corpus of 13 studies, and despite any two of these meta-analyses overlapping with each other between 54% (Eckert et al., 2016 and Linkersdörfer et al., 2012) to 75% (Eckert et al., 2016 and Richlan et al., 2012), not a single resulting region appears in all three meta-analyses.

The lack of strong convergence prompted scrutiny of the original studies (Ramus et al., 2018). Consistent with other earlier publications, this work employed small sample sizes of less than 20 participants in the RD group, and various statistical thresholds, many of which are now considered to be too lenient (Eckert et al., 2016; Ramus et al., 2018). Almost all studies were conducted using voxel-based morphometry (VBM) in Statistical Parametric Mapping (SPM), and there have been no attempts to investigate reproducibility of results using other analysis programs. Most of these studies are based on adults (given the challenges of studying children) and thus likely conflated innate neuroanatomical differences in dyslexia with those consequential to a lifetime of poor reading. While for the most part these studies matched the control and RD groups on age and IQ, few matched the groups on socioeconomic status (SES), which is known to be related to GMV (Lotze et al., 2020; Rakesh et al., 2021) and therefore may have unknowingly driven brain structural differences in dyslexia. Similarly, very few of the studies on RD to date entered any of these as demographic variables as covariates of no interest in the GMV analysis, with none having controlled for age, sex, and SES simultaneously. Further, many studies did not account for global GMV (total GMV) in the analysis in order to differentiate local from global effects. Specifically, only seven of the 13 studies in the meta-analyses entered total GMV (tGMV) as a covariate of no interest, with one correcting for total intracranial volume (tICV) and the other five not correcting for brain size at all. All of these factors likely contributed to the variable results in this corpus of work (Eckert et al., 2016) and, as such, there is concern that the GMV differences reported for RD in the literature are not robust or replicable (Ramus et al., 2018). As in other medical and psychological research, replication studies are challenging due to a lack of details in the published research protocols (Iqbal et al., 2016), prompting the need for a direct replication study.

Here, we leveraged The Adolescent Brain Cognitive Development (ABCD) Study, a demographically diverse sample that is representative of the United States population, to test for GMV differences in large groups with and without RD. The ABCD Study recruited and acquired data in around 11,875 children ages 9–10 years under the same general research protocol (Garavan et al., 2018; Karcher & Barch, 2021). This made it possible to study a large sample of a narrow age range, match the groups on key variables, and also control for these variables in the analyses. We first conducted a direct discovery and replication study, followed by several additional analyses to gauge reproducibility of the results.

For the Discovery and Replication Study, we conducted between-group comparisons of Control versus RD Groups, first for the Discovery Dataset and then again for the Replication Dataset. Both datasets were created to include around 65 participants in each of the four groups, following the recommendation by Ramus and colleagues (2018), who argued that groups below 64 participants are statistically underpowered. We ensured that the Control and RD Groups within each dataset (Discovery and Replication), as well as the four groups overall, were matched on the key demographic variables of age, sex and SES as well as nonverbal reasoning. In addition, given that these variables are known to covary with GMV, they were entered as covariates of no interest in the analyses together with total GMV as recommended by Ramus et al., (2018). We used a voxel-level height threshold of p < 0.005 with a cluster-level threshold of p < 0.05 false discovery rate (FDR) corrected, a more conservative threshold than all but one used in the 13 studies included in the meta-analyses. While this threshold is more stringent than that of this body of studies collectively, some of the more recent studies of RD have adopted an even stricter voxel-level threshold of p < 0.001 (illustrated by six of the 20 studies listed in the supplement in Ramus et al., 2018, as well as a few more since then). We therefore repeated the analyses with this more conservative threshold. Based on the overall findings of the three meta-analyses and a prevalent brain-based model of dyslexia (Pugh et al., 2001; Sandak et al., 2004), we expected the comparison between the Control and RD Groups in both the Discovery and Replication Datasets to reveal relatively less GMV in RD in left-hemisphere perisylvian cortical regions associated with reading (left occipitotemporal, inferior frontal, and temporoparietal cortices).

For the Reproducibility Studies, we conducted several additional analyses to assess factors that influence the results. First, we repeated the Discovery and Replication Study using surface-based morphometry in FreeSurfer instead of VBM in SPM to gauge the effect of an analysis program with a different processing pipeline/software. Second, we combined the two RD and two Control Groups from the Discovery and Replication Datasets to attain a larger sample and used a factorial analysis of variance (ANOVA) to reveal the main effect of Reading Ability (Controls vs. RD, combined across the Discovery and Replication Datasets), the main effect of Dataset (Discovery vs. Replication Dataset, combined across the Control and RD Groups), and their interaction on GMV, the latter quantitatively identifying GMV differences in RD that fail to replicate. For these analyses, we used the same approach as in the initial analysis, as this resulted in larger maps (i.e., maps generated with VBM in SPM using a voxel-level height threshold of p < 0.005 with a cluster-level threshold of p < 0.05 FDR corrected, with age, sex, socioeconomic status, as well as nonverbal reasoning and total GMV as covariates of no interest). Third, to address the effect of accounting for demographic and brain size variables, the ANOVA was repeated after (i) omitting all covariates except for tGMV, to gauge the effect of not controlling for the demographic and nonverbal reasoning variables on the results, (ii) switching out tGMV for tICV, to gauge the effect of not using the same global measure as the variable of interest (i.e., GMV), but an estimate of total brain volume (i.e., tICV) on the results, and (iii) omitting all covariates of no interest, allowing us to gauge the effects of controlling for tGMV (or tICV) on the results. We removed the demographic variables first as this made it possible to test for the outcome of controlling for tGMV in the absence of other variables that likely share variance with brain size.

In sum, in a sample of 262 children, we tested (1) GMV differences between those with and without reading disability using a discovery and a replication approach followed by (2) reproducibility across different methods (analysis program, larger sample ANOVA and covariates of no interest). The results will shed light on how robust GMV differences are in RD and the impact of specific methodological approaches on the findings. Importantly, they will provide a lens by which to view past publications of GMV in dyslexia while also assessing the utility of this neuroanatomical metric going forward.

## 2 Methods

### 2.1 Participant Selection

Participants from the first “baseline” time point of the ABCD Study (n = 11,878) were excluded if they scored below the average range on nonverbal reasoning, ≤ 7 on the Matrix Reasoning subtest of the Wechsler Intelligence Scale for Children-V (roughly equivalent to a standard score of 85) and if they scored ≤ 70 on the NIH Toolbox Picture Vocabulary Test (Gershon et al., 2013, 2014). The remaining participants (n = 9,067) were categorized as potential Control or RD participants based on their NIH Toolbox Oral Reading Recognition Test® score (TORRT; Gershon et al., 2013, 2014): ≥90 as Controls (n = 7,573) and <85 as RD (n = 677). This means the reading proficiency of the RD participants was at least one standard deviation (15 points) below the normative mean (standard score of 100), corresponding to below the 16^th^ percentile, consistent with the approach taken by many prior studies of GMV in RD, so that our results can be interpreted in the context of the literature. We then ruled out those with significant use of another language or non-native users of English by selecting only those who responded “No” when asked “Besides English, do you speak or understand another language or dialect?” on the Youth Acculturation Survey (YAS) Modified from PhenX, combined with the parent reporting that the child’s native language at birth was English. This left 4,452 Control participants and 400 RD participants. We selected RD participants for the Discovery Dataset from five ABCD Study sites and propensity-matched (Kline & Luo, 2022) to Control participants from these same sites on SES (combined household income) and nonverbal reasoning (Matrix Reasoning). SES is total household income (binned) from the Longitudinal Parent Demographics Survey used in the ABCD Study. After conducting MRI quality control (as described below), we arrived at 65 Control and 66 RD participants for the Discovery Dataset. We then selected RD participants for the Replication Dataset from five other ABCD Study sites and here, too, propensity-matched to Control participants from these same sites on SES and nonverbal reasoning. After MRI quality control, we removed two participants randomly to attain 65 Control and 66 RD participants for the Replication Dataset (see Table 1 for demographics and performance for both datasets).

**Table 1.** Participant Demographics.

|  | Discovery Dataset |  |  | Replications Datasets |  |  | Combined Stats |
| --- | --- | --- | --- | --- | --- | --- | --- |
|  | Control | RD | Stats | Control | RD | Stats |  |
| Age | 9.86 ± 0.62 | 9.91 ± 0.64 | $t(129) = -0.42$,<br>$p = 0.674$ | 9.94 ± 0.63 | 10.0 ± 0.62 | $t(129) = -0.61$,<br>$p = 0.542$ | $F(3) = 0.605$,<br>$p = 0.612$ |
| Sex (F/M) | 32/33 | 36/30 | $\chi^2(1) = 0.37$,<br>$p = 0.543$ | 32/33 | 25/41 | $\chi^2(1) = 1.72$,<br>$p = 0.190$ | $\chi^2(3) = 3.91$,<br>$p = 0.271$ |
| SES | 4.48 ± 2.01 | 4.95 ± 2.32 | $t(117) = -1.16$,<br>$p = 0.248$ | 4.61 ± 2.03 | 4.93 ± 2.28 | $t(117) = -0.8$,<br>$p = 0.423$ | $F(3) = 0.681$,<br>$p = 0.564$ |
| Nonverbal Reasoning | 10.6 ± 2.25 | 10.1 ± 1.9 | $t(129) = 1.4$,<br>$p = 0.165$ | 10.6 ± 2.29 | 10.0 ± 1.79 | $t(129) = 1.76$,<br>$p = 0.081$ | $F(3) = 1.68$,<br>$p = 0.172$ |
| Reading | 105 ± 15.7 | 77.2 ± 3.67 | $t(129) = 14.16$,<br>$p < 0.001$ | 106 ± 15.5 | 77.8 ± 3.64 | $t(129) = 14.3$,<br>$p < 0.001$ | $F(3) = 135$,<br>$p < 0.001$ |
| Vocabulary | 107 ± 13.3 | 96.2 ± 10.9 | $t(129) = 4.914$,<br>$p < 0.001$ | 103 ± 15.9 | 91.3 ± 11.4 | $t(129) = 4.88$,<br>$p < 0.001$ | $F(3) = 18.3$,<br>$p < 0.001$ |

Note: Discovery and Replication Datasets. Counts reported for sex (F/M), and averages and standard deviations reported for Age, SES, Nonverbal Reasoning, Reading, and Vocabulary. T- tests and $\chi^2$ tests for the two groups in the Discovery Dataset are reported in the fourth column, for the two groups in the Replication Dataset in the seventh column, and a one-way ANOVA for all four groups in the last column. SES (Socioeconomic Status) = Total Household Income bin from the Longitudinal Parent Demographics Survey; Nonverbal Reasoning = Matrix Reasoning Test; Reading = NIH Toolbox Oral Reading Recognition Test; Vocabulary = NIH Toolbox Picture Vocabulary Test.

### 2.2 MRI Data Acquisition

T1-weighted MR images with a 1 x 1 x 1 mm^3^ voxel resolution were acquired on 3T MR scanners. For more information on the MR scanners and parameters, see https://abcdstudy.org/images/Protocol_Imaging_Sequences.pdf.

### 2.3 Discovery and Replication

#### 2.3.1 GMV Analysis using Voxel-Based Morphometry (VBM) in Statistical Parametric Mapping (SPM)

##### 2.3.1.1 MRI Data Quality Control and Preprocessing for VBM in SPM

For the GMV analysis using VBM in SPM (Wellcome Trust Centre for Neuroimaging; http://www.fil.ion.ucl.ac.uk/spm), we applied two quality control steps. First, two researchers blinded to group membership assessed the MR images and excluded those images (a) with anatomical anomalies and (b) with a score of 1 or 2 on an image quality scale of 1 (severely distorted) to 5 (optimal). Second, we used the Image Quality Rating (IQR) reports from the Computational Anatomy Toolbox 12 (https://neuro-jena.github.io/cat12-help/), in SPM12. The reports quantify the amount of noise and distortion and we excluded images with ratings below 70% from the analyses. Nine potential RD and eleven potential Control participants from the Discovery Dataset, and four potential RD and nine potential Control participants from the Replication Dataset did not meet these MRI quality control criteria and were excluded.

Preprocessing of all images in VBM (Ashburner & Friston, 2000) via SPM12 was performed as outlined by Ashburner (2015). The images were co-registered to the white matter tissue probability map and then segmented into gray matter, white matter, and cerebrospinal fluid images. Images were used to create study-specific templates for each analysis and spatially normalized to the Montreal Neurological Institute (MNI) stereotaxic space via affine registration of the generated template to the MNI template using DARTEL (Ashburner, 2007). Images were examined to confirm successful normalization, smoothed with a Gaussian kernel of 8 mm full width at half maximum (FWHM) and intensity thresholding was set to 0.2.

##### 2.3.1.2 Analysis for VBM in SPM: Differences in GMV between Control and RD Groups in Discovery and Replication Study (two thresholds)

We compared GMV using VBM between the RD and Control Groups in the Discovery Dataset and then again in the Replication Dataset using two-sample t-tests, applying a voxel-wise height threshold of p < 0.005 and cluster-level extent threshold of p < 0.05, FDR, with age, sex, SES, study site, nonverbal reasoning and tGMV entered as covariates of no interest. While this threshold is more stringent than those use in almost all of the 13 studies in the meta-analyses, we also repeated the two-sample t-tests using an even more stringent threshold of voxel-wise height threshold of p < 0.001 and a cluster-level extent threshold of p < 0.05, FDR to reduce false positive rates (Woo et al., 2014).

### 2.4 Reproducibility Studies

#### 2.4.1 GMV Analysis using Surface-Based Morphometry (SBM) in FreeSurfer

##### 2.4.1.1 MRI Data Quality Control and Preprocessing for SBM in FreeSurfer

For the FreeSurfer analysis, structural MRI data underwent quality control and were processed using the standardized ABCD Study image processing pipeline implemented by the Data Analysis, Informatics, and Resource Center (DAIRC) as described in Hagler et al. (2019). This pipeline includes correction for gradient nonlinearity distortions and intensity inhomogeneity correction. Cortical reconstruction and segmentation were performed using the FreeSurfer pipeline (Fischl, 2012), which includes skull stripping, white matter segmentation, surface reconstruction, correction of topological defects, and surface optimization (Dale et al., 1999; Fischl et al., 2001). Cortical surfaces were registered to the Destrieux Atlas (Destrieux et al., 2010), from which regional gray matter volume measures were extracted.

##### 2.4.1.2 Analysis for SBM in FreeSurfer: Differences in GMV between Control and RD Groups in Discovery and Replication Study

Here we again compared GMV between the RD and Control Groups in the Discovery Dataset and then again in the Replication Dataset this time using FreeSurfer. Between-group comparisons were conducted using one-way analyses of covariance (ANCOVAs) for each of the 148 regions of the Destrieux Atlas. All analyses were conducted in R, with significance set at *p* < .05 with FDR correction for the 148 regions and covariates of no interest were the same as those included in the VBM analyses.

#### 2.4.2 Analysis for VBM in SPM: 2x2 ANOVA when Accounting for Different Variable

Using the preprocessed VBM data from above (see 2.2.1.2) that had been used for the t-tests, a factorial design was used to combine all four groups to test for the main effect of Reading Ability (Control vs. RD while combining across both datasets), main effect of Dataset (Discovery vs. Replication while combining across the groups of different reading abilities), and their interaction. The statistical approach was consistent with the main VBM between-group comparison above (voxel-wise height threshold of p < 0.005 and cluster-level extent threshold of p < 0.05, FDR, with age, sex, SES, study site, nonverbal reasoning and tGMV entered as covariates of no interest). To gauge the role of these variables, we repeated the ANOVA but included only tGMV as a covariate of no interest, allowing us to assess the role of age, sex, SES, study site, and nonverbal reasoning on the results. Next, to gauge the role of tGMV, the ANOVA was repeated, but this time only controlling for total intracranial volume (tICV) or not controlling for any covariates. The demographic and performance variables were not included in the last two analyses as they are related to brain size and would therefore obscure the impact of controlling for tGMV or tICV.

For all VBM in SPM analyses, Brodmann areas (BA) and anatomical labels of the coordinates from SPM12 were determined using the label4MRI package (https://github.com/yunshiuan/label4MRI). This package uses the atlas of Brodmann areas and the Automated Anatomical Labeling Atlas (Tzourio-Mazoyer et al., 2002). Each peak was limited to a maximum of two subpeaks as per the SPM12 default settings.

## 3 Results

### 3.1 Participant Profiles

As can be seen in Table 1, in the Discovery Dataset, as well as in the Replication Dataset, the respective Control and RD Groups did not differ significantly on age, sex distribution, SES (combined household income), or nonverbal reasoning (Matrix Reasoning), as expected. Also, as designed, the respective RD Groups had significantly lower reading scores (TORRT) than the Control Groups and, as expected, significantly lower vocabulary scores (Picture Vocabulary). Notably, the group mean performance for reading was almost two standard deviations lower in the RD Groups than the respective Control Groups in both the Discovery (difference of 27.8 standard score points) and Replication (difference of 28.2 standard score points) Dataset, indicating a robust impediment in the RD Groups relative to the Controls.

As expected, a one-way ANOVA comparing all four groups (Discovery and the Replication Datasets) revealed no significant difference in age, sex, SES, or nonverbal reasoning while reading (and vocabulary) scores were significantly different due to the much lower reading ability of the RD Groups (Table 1).

MRI data analysis (VBM in SPM) yielded participants’ tGMV, an important variable to account for (together with the demographic variables noted here). A one-way ANOVA revealed significant differences in the means of the four groups (F(3) = 3.57, p = 0.015), with the Control Groups having lower tGMV than the RD Groups in the Discovery (t(129) = 3.16, p < 0.033) and the Replication (t(129) = 2.36, p < 0.020) Datasets.

### 3.2 MRI Results

#### 3.2.1 Discovery and Replication Study

##### 3.2.1.1 Results for VBM in SPM: Differences in GMV between Control and RD Groups in Discovery and Replication Study (two thresholds)

We first tested for GMV differences between the Control and RD Groups using VBM in th Discovery Dataset and then in the Replication Dataset. For the Discovery Dataset at a voxel-wise height threshold of p < 0.005 (Figure 1A, Table 2 A), the Control Group had more GMV than the RD Group in the left inferior temporal gyrus (BA 20), left cerebellum lobule VI, and right superior temporal gyrus (BA 22). On the other hand, the Control Group had less GMV than the RD Group in the right middle orbitofrontal gyrus (BA 10). When repeating the analysis with a voxel-wise height threshold of p < 0.001 (Table 2 C), the Control Group had more GMV than th RD Group in the left calcarine fissure (BA 17), and there were no regions where the Control Group had less GMV than the RD Group.

**Figure 1.**
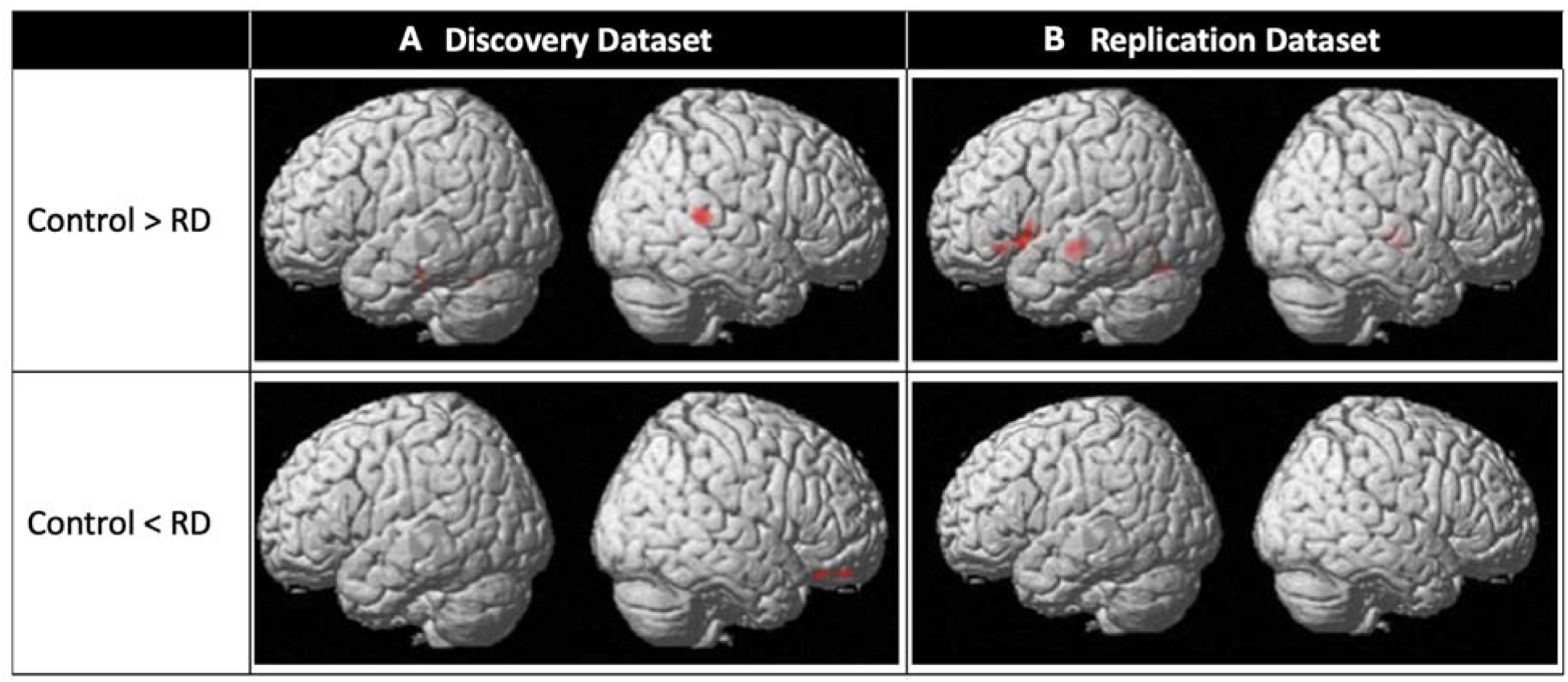
Differences in GMV between the Control Groups and the RD Groups in the (A) Discovery Dataset and (B) Replication Dataset. Maps were generated in VBM in SPM using a two-sample t-test (voxel-wise height threshold of p < 0.005; cluster-level extent threshold of p < 0.05, FDR). The corresponding coordinates for significant clusters are in Table 2. None of the between-group differences in RD identified in each dataset are shared across datasets.

**Table 2.** Differences in GMV (VBM in SPM) between the Control Groups and the RD Groups in Discovery Dataset (A and C), and Replication Dataset (B and D) at voxel-wise height thresholds p < 0.005 and p < 0.001.

| **A. Discovery Dataset** |  |  |  |  |  |  |
| --- | --- | --- | --- | --- | --- | --- |
| **x** | **y** | **z** | **Anatomical Region** | **BA** | **Z** | **kE** |
| **Control > RD ($p < 0.005$)** |  |  |  |  |  |  |
| -46 | -21 | -20 | Left Inferior Temporal Gyrus | 20 | 3.3 | 234 |
| -45 | -27 | -33 | Left Inferior Temporal Gyrus | 20 | 3.17 |  |
| -38 | -26 | -20 | Left Fusiform Gyrus | 36 | 3.01 |  |
| -14 | -62 | -27 | Left Cerebellum Lobule VI |  | 3.63 | 344 |
| 54 | -36 | 10 | Right Superior Temporal Gyrus | 22 | 3.62 | 547 |
| 45 | -42 | 3 | Right Middle Temporal Gyrus | 21 | 3.09 |  |
| **Control < RD ($p < 0.005$)** |  |  |  |  |  |  |
| 32 | 50 | -21 | Right Middle Orbitofrontal Gyrus | 10 | 3.87 | 328 |
| 21 | 48 | -24 | Right Middle Orbitofrontal Gyrus | 11 | 3.7 |  |
| 33 | 36 | -24 | Right Inferior Orbitofrontal Gyrus | 47 | 3.09 |  |
| **B. Replication Dataset** |  |  |  |  |  |  |
| **x** | **y** | **z** | **Anatomical Region** | **BA** | **Z** | **kE** |
| **Control > RD ($p < 0.005$)** |  |  |  |  |  |  |
| -38 | 28 | -9 | Left Inferior Orbitofrontal Gyrus | 47 | 3.85 | 461 |
| -44 | 15 | -6 | Left Insula | 47 | 3.12 |  |
| -44 | 4 | 0 | Left Insula | 13 | 2.78 |  |
| -21 | 21 | -3 | Left Putamen | 49 | 3.36 | 341 |
| -27 | 2 | 14 | Left Putamen | 49 | 2.97 |  |
| -48 | -15 | -6 | Left Middle Temporal Gyrus | 22 | 3.38 | 470 |
| -48 | -20 | -16 | Left Middle Temporal Gyrus | 21 | 3.05 |  |
| -16 | -45 | -9 | Left Lingual Gyrus | 19 | 3.54 | 1182 |
| -21 | -63 | -3 | Left Lingual Gyrus | 19 | 3.32 |  |
| -16 | -64 | -10 | Left Lingual Gyrus | 19 | 3.08 |  |
| 45 | -15 | -4 | Right Superior Temporal Gyrus | 22 | 3.26 | 432 |
| 42 | -24 | 0 | Right Superior Temporal Gyrus | 41 | 3.25 |  |
| 42 | -20 | -10 | Right Hippocampus | 22 | 3.19 |  |
| **Control < RD ($p < 0.005$)** |  |  |  |  |  |  |
|  |  |  | None |  |  |  |
| **C. Discovery Dataset** |  |  |  |  |  |  |
| **x** | **y** | **z** | **Anatomical Region** | **BA** | **Z** | **kE** |
| **Control > RD ($p < 0.001$)** |  |  |  |  |  |  |
| -11 | -89 | 6 | Left Calcarine Fissure | 17 | 4.14 | 83 |

| Control < RD (p < 0.001) |  |  |  |  |  |  |
| --- | --- | --- | --- | --- | --- | --- |
|  |  |  | None |  |  |  |
| D. Replication Dataset |  |  |  |  |  |  |
| x | y | z | Anatomical Region | BA | Z | kE |
| Control > RD (p < 0.001) |  |  |  |  |  |  |
|  |  |  | None |  |  |  |
| Control < RD (p < 0.001) |  |  |  |  |  |  |
|  |  |  | None |  |  |  |

*Note: Black font indicates peak coordinates of cluster and its cluster size, and gray font indicates sub-peak coordinates within that cluster. Coordinates indicating anatomical location are in Montreal Neurological Institute (MNI) stereotaxic space. For results A and B, see Figure 1.*

For the Replication Dataset at voxel-wise height threshold of p < 0.005 (Figure 1B, Table 2 B), the Control Group had more GMV than the RD Group in the left inferior orbitofrontal gyrus (BA 47), left putamen (BA 49), left middle temporal gyrus (BA 22), left lingual gyrus (BA 19) and two foci in the right superior temporal gyrus (BA 22 and BA 41). There were no findings of less GMV in the Control Group relative to the RD Group. Of note, right superior temporal gyru (STG) is reported for both datasets, but the locations within the STG did not overlap (with the replication finding location more anteriorly and inferiorly than that of the Discovery Dataset).

When repeating the analysis with a voxel-wise height threshold of p < 0.001 (Table 2D), there were no regions where the Control Group had more GMV than the RD Group or vice versa.

#### 3.2.2 Reproducibility Studies

##### 3.2.2.1 Results for SBM in FreeSurfer: Differences in GMV between Control and RD Groups in Discovery and Replication Study

We then tested for GMV differences between the Control and RD Groups (in the Discovery Dataset and then in the Replication Dataset) again, this time using FreeSurfer for the analysis. For the Discovery Dataset (Table 3 A), the Control Group had more GMV than the RD Group in the left anterior transverse collateral sulcus. For the contrast of less GMV in the Control Group than the RD Group there were no findings.

**Table 3.**
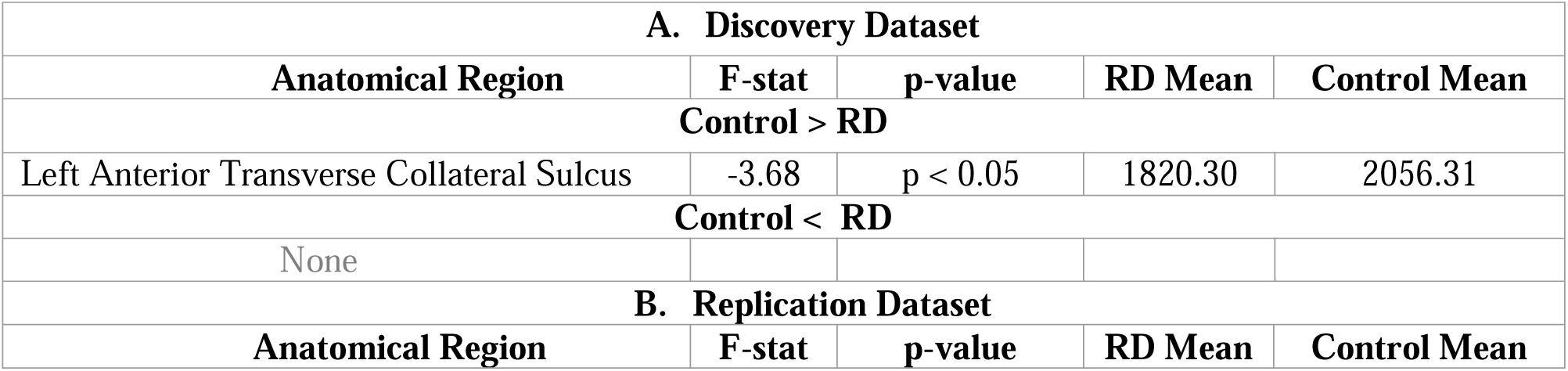
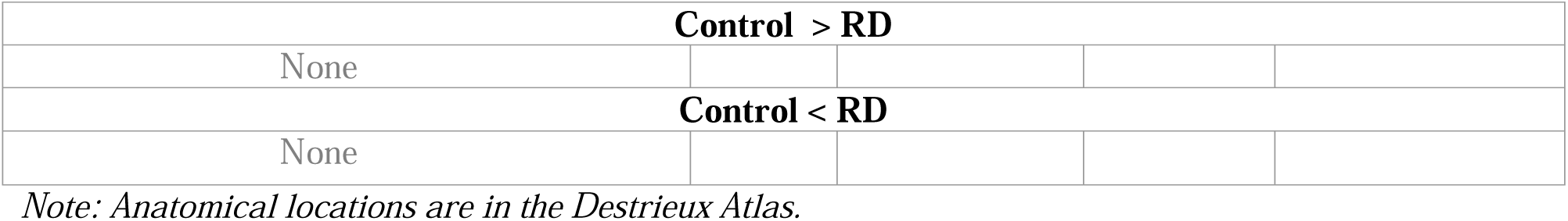
Differences in GMV (FreeSurfer) between the Control Groups and the RD Groups in Discovery Dataset (A), and Replication Dataset (B)

| A. Discovery Dataset |  |  |  |  |
| --- | --- | --- | --- | --- |
| Anatomical Region | F-stat | p-value | RD Mean | Control Mean |
| Control > RD |  |  |  |  |
| Left Anterior Transverse Collateral Sulcus | -3.68 | p < 0.05 | 1820.30 | 2056.31 |
| Control < RD |  |  |  |  |
| None |  |  |  |  |
| B. Replication Dataset |  |  |  |  |
| Anatomical Region | F-stat | p-value | RD Mean | Control Mean |

| **Control > RD** |
| --- |
| None |
| **Control < RD** |
| None |

*Note: Anatomical locations are in the Destrieux Atlas.*

Using FreeSurfer for the analysis of the Replication Dataset (Table 3 B), there were no regions where the Control Group had more GMV than the RD Group or vice versa.

##### 3.2.2.2 Results for VBM in SPM: 2x2 ANOVA

Using VBM in SPM (see 2.4.2), a factorial analysis was used to combine all four groups and test for the main effect of Reading Ability, main effect of Dataset, and their interaction (voxel-wise height threshold of p < 0.005, age, sex and socioeconomic status, study site, nonverbal reasoning and tGMV entered as covariates of no interest). The ANOVA was then repeated three more times, including only tGMV, then including only tICV, and then without covariates of no interest.

###### 3.2.2.2.1 ANOVA: Accounting for Age, Sex, SES, Study Site, Nonverbal Reasoning, and Total GMV

The main ANOVA with age, sex, SES, study site, nonverbal reasoning, and tGMV as covariates of no interest (voxel-wise height threshold of p < 0.005) revealed a main effect of Reading Ability on GMV in six regions (Figure 2 first column, and Table 4). In five of these, the Control Group (Discovery and Replication Datasets combined) had more GMV than the RD Group (Discovery and Replication Datasets combined) and were located in left fusiform gyrus (BA 37) and left cerebellar lobule VI; and in right caudate (BA 48), right superior temporal gyrus (BA 22), and right cerebellar lobule VI. In one region the Control Group had less GMV than the RD Group and it was in the right middle orbitofrontal gyrus (BA 10). As expected, some of these results align with the above results from the Discovery Dataset (e.g., left cerebellar lobule VI), the Replication Dataset (e.g., putamen), or are novel (e.g., right cerebellum lobule VI).

**Figure 2.**
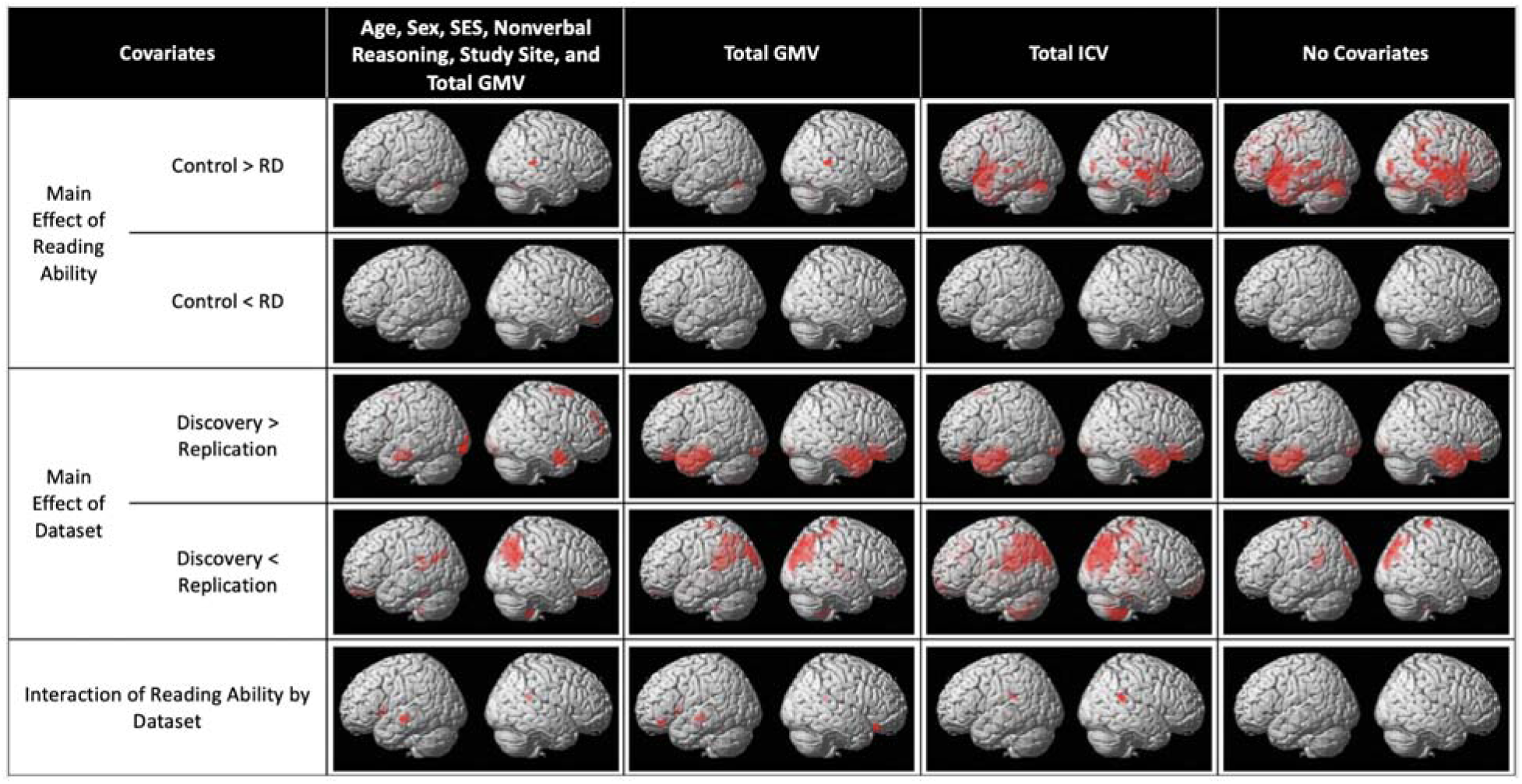
Reproducibility Study (VBM in SPM): ANOVAs to test for Main Effect of Reading Ability, Main Effect of Dataset and their Interactions and three further ANOVAs using different choices of covariates. Maps were generated using a voxel-wise height threshold p < 0.005, cluster-level extent threshold p < 0.05, FDR. First column depicts the main ANOVA result, controlling for Age, Sex, SES, Nonverbal Reasoning, Study Site, and Total GMV. Remaining ANOVA results when these demographic and performance variables are not controlled for, instead only controlling for either total GMV (second column), or total intracranial volume (third column), or using no covariates (last column). Generally, analyses controlling for fewer variables of no interest resulted in more extensive GMV differences between the Control and RD Groups. The corresponding coordinates for significant clusters for the main ANOVA are in Table 4, and for the remaining ANOVAs in Supplemental Table 1, 2 and 3. Figure 3 depicts the interaction result of the main ANOVA.

**Table 4.**
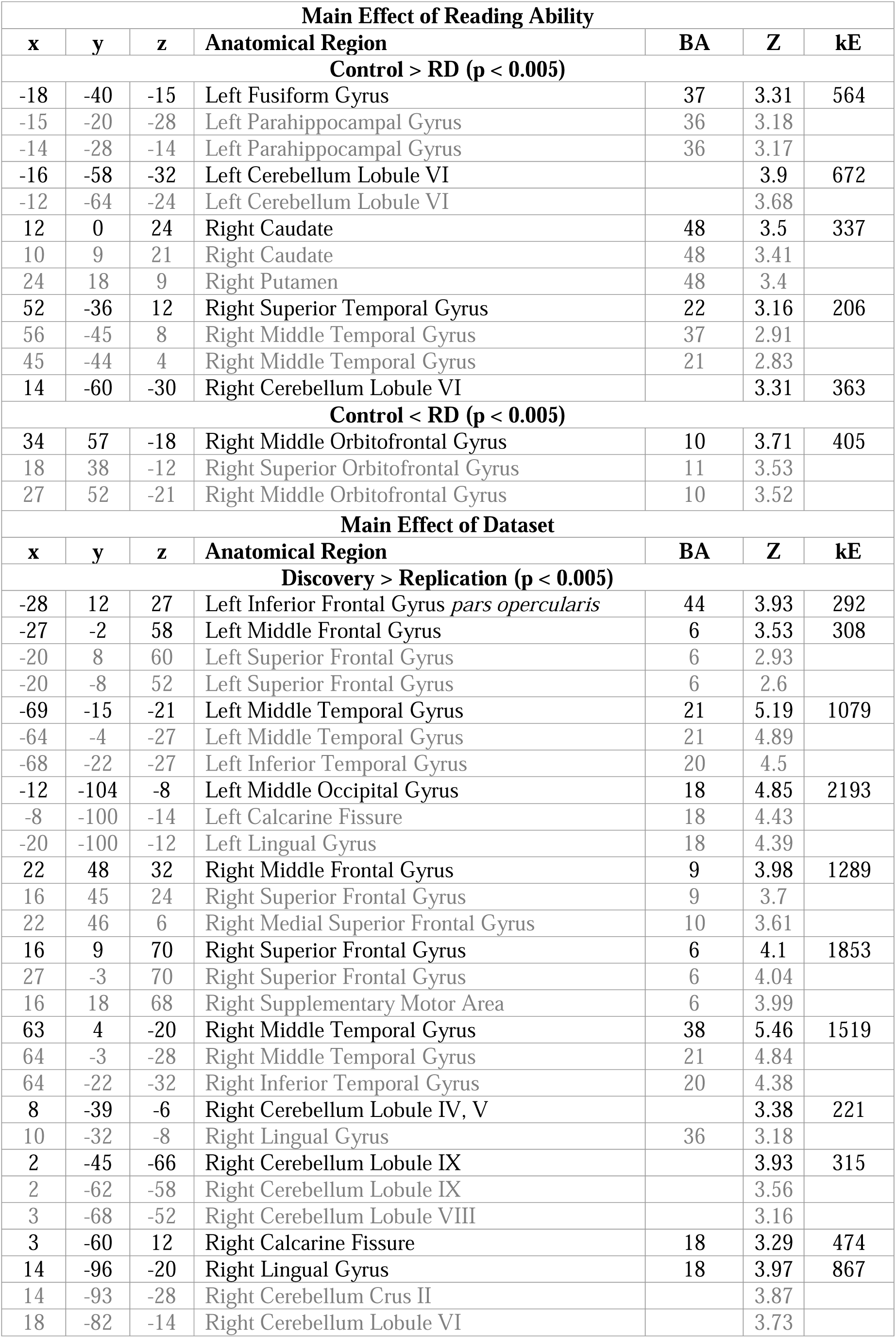
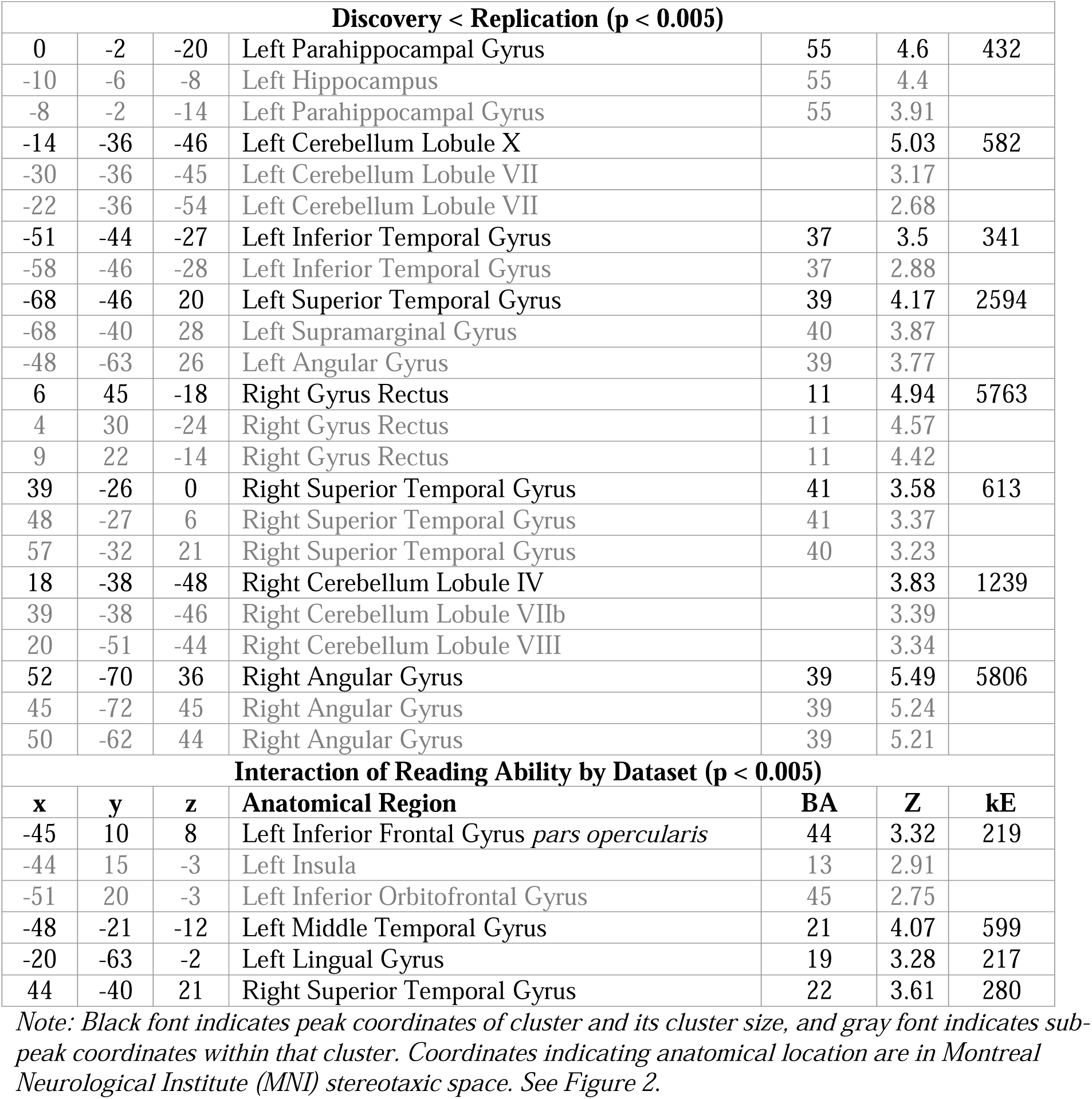
ANOVA (VBM in SPM): Main Effect of Reading Ability, Main Effect of Dataset, and their Interactions with Age, Sex, SES, Study Site, Nonverbal Reasoning, and Total GMV as Covariates.

| x | y | z | Anatomical Region | BA | Z | kE |
| --- | --- | --- | --- | --- | --- | --- |
| **Control > RD (p < 0.005)** |  |  |  |  |  |  |
| -18 | -40 | -15 | Left Fusiform Gyrus | 37 | 3.31 | 564 |
| -15 | -20 | -28 | Left Parahippocampal Gyrus | 36 | 3.18 |  |
| -14 | -28 | -14 | Left Parahippocampal Gyrus | 36 | 3.17 |  |
| -16 | -58 | -32 | Left Cerebellum Lobule VI |  | 3.9 | 672 |
| -12 | -64 | -24 | Left Cerebellum Lobule VI |  | 3.68 |  |
| 12 | 0 | 24 | Right Caudate | 48 | 3.5 | 337 |
| 10 | 9 | 21 | Right Caudate | 48 | 3.41 |  |
| 24 | 18 | 9 | Right Putamen | 48 | 3.4 |  |
| 52 | -36 | 12 | Right Superior Temporal Gyrus | 22 | 3.16 | 206 |
| 56 | -45 | 8 | Right Middle Temporal Gyrus | 37 | 2.91 |  |
| 45 | -44 | 4 | Right Middle Temporal Gyrus | 21 | 2.83 |  |
| 14 | -60 | -30 | Right Cerebellum Lobule VI |  | 3.31 | 363 |
| **Control < RD (p < 0.005)** |  |  |  |  |  |  |
| 34 | 57 | -18 | Right Middle Orbitofrontal Gyrus | 10 | 3.71 | 405 |
| 18 | 38 | -12 | Right Superior Orbitofrontal Gyrus | 11 | 3.53 |  |
| 27 | 52 | -21 | Right Middle Orbitofrontal Gyrus | 10 | 3.52 |  |
| **Main Effect of Dataset** |  |  |  |  |  |  |
| x | y | z | Anatomical Region | BA | Z | kE |
| **Discovery > Replication (p < 0.005)** |  |  |  |  |  |  |
| -28 | 12 | 27 | Left Inferior Frontal Gyrus *pars opercularis* | 44 | 3.93 | 292 |
| -27 | -2 | 58 | Left Middle Frontal Gyrus | 6 | 3.53 | 308 |
| -20 | 8 | 60 | Left Superior Frontal Gyrus | 6 | 2.93 |  |
| -20 | -8 | 52 | Left Superior Frontal Gyrus | 6 | 2.6 |  |
| -69 | -15 | -21 | Left Middle Temporal Gyrus | 21 | 5.19 | 1079 |
| -64 | -4 | -27 | Left Middle Temporal Gyrus | 21 | 4.89 |  |
| -68 | -22 | -27 | Left Inferior Temporal Gyrus | 20 | 4.5 |  |
| -12 | -104 | -8 | Left Middle Occipital Gyrus | 18 | 4.85 | 2193 |
| -8 | -100 | -14 | Left Calcarine Fissure | 18 | 4.43 |  |
| -20 | -100 | -12 | Left Lingual Gyrus | 18 | 4.39 |  |
| 22 | 48 | 32 | Right Middle Frontal Gyrus | 9 | 3.98 | 1289 |
| 16 | 45 | 24 | Right Superior Frontal Gyrus | 9 | 3.7 |  |
| 22 | 46 | 6 | Right Medial Superior Frontal Gyrus | 10 | 3.61 |  |
| 16 | 9 | 70 | Right Superior Frontal Gyrus | 6 | 4.1 | 1853 |
| 27 | -3 | 70 | Right Superior Frontal Gyrus | 6 | 4.04 |  |
| 16 | 18 | 68 | Right Supplementary Motor Area | 6 | 3.99 |  |
| 63 | 4 | -20 | Right Middle Temporal Gyrus | 38 | 5.46 | 1519 |
| 64 | -3 | -28 | Right Middle Temporal Gyrus | 21 | 4.84 |  |
| 64 | -22 | -32 | Right Inferior Temporal Gyrus | 20 | 4.38 |  |
| 8 | -39 | -6 | Right Cerebellum Lobule IV, V |  | 3.38 | 221 |
| 10 | -32 | -8 | Right Lingual Gyrus | 36 | 3.18 |  |
| 2 | -45 | -66 | Right Cerebellum Lobule IX |  | 3.93 | 315 |
| 2 | -62 | -58 | Right Cerebellum Lobule IX |  | 3.56 |  |
| 3 | -68 | -52 | Right Cerebellum Lobule VIII |  | 3.16 |  |
| 3 | -60 | 12 | Right Calcarine Fissure | 18 | 3.29 | 474 |
| 14 | -96 | -20 | Right Lingual Gyrus | 18 | 3.97 | 867 |
| 14 | -93 | -28 | Right Cerebellum Crus II |  | 3.87 |  |
| 18 | -82 | -14 | Right Cerebellum Lobule VI |  | 3.73 |  |

| Discovery < Replication (p < 0.005) |  |  |  |  |  |  |
| --- | --- | --- | --- | --- | --- | --- |
| 0 | -2 | -20 | Left Parahippocampal Gyrus | 55 | 4.6 | 432 |
| -10 | -6 | -8 | Left Hippocampus | 55 | 4.4 |  |
| -8 | -2 | -14 | Left Parahippocampal Gyrus | 55 | 3.91 |  |
| -14 | -36 | -46 | Left Cerebellum Lobule X |  | 5.03 | 582 |
| -30 | -36 | -45 | Left Cerebellum Lobule VII |  | 3.17 |  |
| -22 | -36 | -54 | Left Cerebellum Lobule VII |  | 2.68 |  |
| -51 | -44 | -27 | Left Inferior Temporal Gyrus | 37 | 3.5 | 341 |
| -58 | -46 | -28 | Left Inferior Temporal Gyrus | 37 | 2.88 |  |
| -68 | -46 | 20 | Left Superior Temporal Gyrus | 39 | 4.17 | 2594 |
| -68 | -40 | 28 | Left Supramarginal Gyrus | 40 | 3.87 |  |
| -48 | -63 | 26 | Left Angular Gyrus | 39 | 3.77 |  |
| 6 | 45 | -18 | Right Gyrus Rectus | 11 | 4.94 | 5763 |
| 4 | 30 | -24 | Right Gyrus Rectus | 11 | 4.57 |  |
| 9 | 22 | -14 | Right Gyrus Rectus | 11 | 4.42 |  |
| 39 | -26 | 0 | Right Superior Temporal Gyrus | 41 | 3.58 | 613 |
| 48 | -27 | 6 | Right Superior Temporal Gyrus | 41 | 3.37 |  |
| 57 | -32 | 21 | Right Superior Temporal Gyrus | 40 | 3.23 |  |
| 18 | -38 | -48 | Right Cerebellum Lobule IV |  | 3.83 | 1239 |
| 39 | -38 | -46 | Right Cerebellum Lobule VIIb |  | 3.39 |  |
| 20 | -51 | -44 | Right Cerebellum Lobule VIII |  | 3.34 |  |
| 52 | -70 | 36 | Right Angular Gyrus | 39 | 5.49 | 5806 |
| 45 | -72 | 45 | Right Angular Gyrus | 39 | 5.24 |  |
| 50 | -62 | 44 | Right Angular Gyrus | 39 | 5.21 |  |
| Interaction of Reading Ability by Dataset (p < 0.005) |  |  |  |  |  |  |
| x | y | z | Anatomical Region | BA | Z | kE |
| -45 | 10 | 8 | Left Inferior Frontal Gyrus *pars opercularis* | 44 | 3.32 | 219 |
| -44 | 15 | -3 | Left Insula | 13 | 2.91 |  |
| -51 | 20 | -3 | Left Inferior Orbitofrontal Gyrus | 45 | 2.75 |  |
| -48 | -21 | -12 | Left Middle Temporal Gyrus | 21 | 4.07 | 599 |
| -20 | -63 | -2 | Left Lingual Gyrus | 19 | 3.28 | 217 |
| 44 | -40 | 21 | Right Superior Temporal Gyrus | 22 | 3.61 | 280 |

Note: Black font indicates peak coordinates of cluster and its cluster size, and gray font indicates sub-peak coordinates within that cluster. Coordinates indicating anatomical location are in Montreal Neurological Institute (MNI) stereotaxic space. See Figure 2.

The main effect of Dataset on GMV yielded a larger number, with 19 regions (Figure 2 first column, and Table 4). In 11 of these, the Discovery Dataset (Control and RD Groups combined) had greater GMV than the Replication Dataset (Control and RD Groups combined) and these four left-hemisphere and seven right-hemisphere regions are listed in Table 4. Eight regions that had less GMV in the Discovery Dataset than the Replication Dataset, resided in four left and four right hemisphere regions listed in Table 4. Of note are the results in the left inferior frontal gyrus *pars opercularis* (BA 44) (Discovery>Replication Dataset) and left superior temporal gyrus (BA39) (Discovery<Replication Dataset), given that these regions are associated with RD, but here were found to differ between two sets of participants.

Finally, there was an interaction effect on GMV between Reading Ability and Dataset in four regions: left inferior frontal gyrus *pars opercularis* (BA 44), left middle temporal gyrus (BA 21), left lingual gyrus (BA 19), and right superior temporal gyrus (BA 22). Table 4 details their locations, and Figure 3 illustrates the nature of the interactions: in the three left hemisphere regions, the Control Group had more GMV than the RD Group in the Replication Dataset, but this between-group difference did not emerge from the Discovery Dataset, while in the right superior temporal gyrus, the Control Group had more GMV than the RD Group in the Discovery Dataset, but the two groups did not differ in the Replication Dataset. As would be expected, these results directly map onto the three left hemisphere regions (inferior frontal, middle temporal and lingual gyri) emerging only from the Replication Dataset result, and the right superior temporal gyrus emerging only from the Discovery Dataset result, described above.

**Figure 3.**
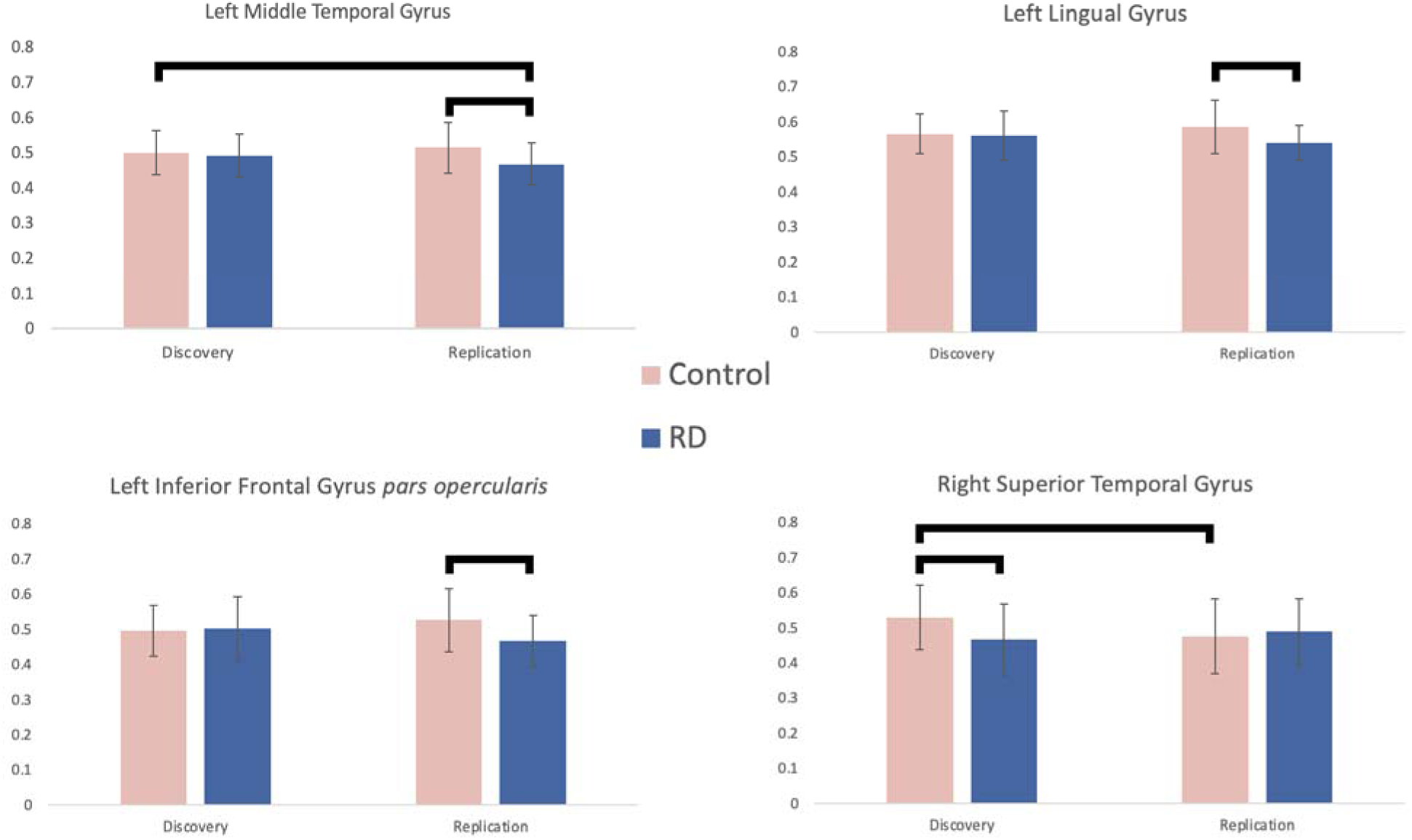
Interaction between Reading Ability and Dataset (VBM in SPM). Age, Sex, SES, Study Site, Nonverbal Reasoning, and Total GMV as Covariates. Whole brain analysis, with voxel-wise height threshold p < 0.005, cluster-level extent threshold p < 0.05, FDR. Brackets above graphs represent significant differences from *post hoc* t-tests (p < 0.05).

###### 3.2.2.2.2 ANOVA: Accounting for Total GMV (tGMV)

This ANOVA was conducted as a repetition of the above ANOVA, but without including age, sex, SES, study site, and nonverbal reasoning, and only controlling for tGMV. The main effect of Reading Ability on GMV revealed four regions (Figure 2, Supplement Table 1). For all of these, the Control Group (Discovery and Replication Datasets combined) had more GMV than the RD Group (Discovery and Replication Datasets combined): left cerebellar lobule VI, right caudat (BA 48), right superior temporal gyrus (BA 22), and right cerebellum lobule VI, which also emerged from the main ANOVA, but this time without left fusiform. Also, this time there was no result for less GMV in Control Group than in the RD Group (i.e., no right middle orbitofrontal gyrus).

There was a main effect of Dataset on GMV in 22 brain regions. For 10 of these, the Discovery Dataset (Control and RD groups combined) had more GMV than the Replication Dataset (Control and RD groups combined), and for 12 regions it was the opposite contrast, as listed in Supplemental Table 1.

Finally, there was an interaction between Reading Ability and Dataset in five regions: left inferior orbitofrontal gyrus (BA 47), left inferior frontal gyrus *pars opercularis* (BA 44), left middle temporal gyrus (BA 22), right inferior orbitofrontal gyrus (BA 47), and right superior temporal gyrus (BA 22), where the difference between Control and RD Groups presented differently depending on the Dataset. Three of these had emerged from the main ANOVA described above (left inferior frontal gyrus *pars opercularis*, left middle temporal gyrus, and right superior temporal gyrus).

###### 3.2.2.2.3 ANOVA: Accounting for Total Intracranial Volume (tICV)

The ANOVA that only included tICV as a covariate of no interest revealed a main effect of Reading Ability on GMV in six regions (Figure 2, Supplement Table 2), and for all of these, the Control Group (Discovery and Replication Datasets combined) had more GMV than the RD Group (Discovery and Replication Datasets combined): left middle frontal gyrus (BA 6), left precentral gyrus (BA 4), left middle temporal gyrus (BA 21), left cerebellum lobule VI, right precentral gyrus (BA 6), and right middle occipital gyrus (BA 19). Notably, only one finding (left cerebellum) overlapped with the results of the main ANOVA above, this time much larger in size and including left fusiform gyrus and right cerebellum, and this time there was also less GMV in RD left temporal and frontal regions. There again was no region that had less GMV in the Control Group than in the RD Group.

For the main effect of Dataset, there were 13 results, of which five were attributed to more GMV in the Discovery Dataset than the Replication Dataset, and eight to the opposite contrast as listed in Supplement Table 2. For both main effects results, the clusters were larger and more widespread than those seen in the main ANOVA.

Finally, the interaction of Reading Ability and Dataset revealed only two regions, in left and right superior temporal gyri (BA 22) where the difference between the Control and RD Groups presented differently depending on the Dataset. The right superior temporal gyrus was the same as observed in the interaction analysis of the main ANOVA.

###### 3.2.2.2.4 ANOVA: no Covariates

For the ANOVA that did not include any covariates (Figure 2, Supplement Table 3), there was a main effect of Reading Ability on GMV in two very large clusters, for both of which GMV was more in the Control Group than the RD Group, with peaks in the right precentral gyrus (BA 6) and extending throughout all lobes of the brain, and in the right cerebellum lobule VI extending to the left cerebellum. There were no regions that had less GMV in the Control Group than the RD Group. The main effect of Dataset revealed 15 regions with differences between the Discovery and Replication Dataset in widespread regions throughout the brain, seven for more GMV in the Discovery Dataset than the Replication Dataset, and eight for the reverse comparison. Both main effects revealed large clusters that were widespread and with fewer foci than in the above-described results. Unlike the above ANOVA results, the interaction of Reading Ability by Dataset revealed no regions.

## 4 Discussion

It is generally accepted that research findings for GMV differences in those with reading disability converge in bilateral temporoparietal and left occipitotemporal and prefrontal cortices as well as the cerebellum (Eckert et al., 2016; Linkersdörfer et al., 2012; Richlan et al., 2012). These aberrations, particularly in left hemisphere perisylvian cortical regions known to be involved in language, align with dyslexia being a language-based learning disability and with a prevailing brain-based model that describes how these regions subserve reading acquisition and deviate in those with dyslexia (Pugh et al., 2001; Sandak et al., 2004). However, concerns have been raised that consistency among these GMV findings in RD is in fact low (Ramus et al., 2018) and the results may not be robust (Eckert et al., 2018). Lack of convergence has been attributed to a number of shortcomings generally found in older neuroimaging studies: small sample sizes, lenient statistical thresholds and insufficient accounting for demographic variables and global GMV.

In the current study, we addressed these concerns using a direct Discovery and Replication Study, using larger group sizes and more conservative statistical thresholds than those generally found in the GMV RD literature, as well as matching the groups on critical variables, as well as controlling for these variables in the analyses. We found that between-group differences in GMV did not replicate across the two datasets.

Next, we addressed these concerns with Reproducibility Studies and first repeated the comparisons of GMV between the same Control and RD Groups in both datasets (Discovery and Replication) using a different analysis program for measuring GMV. We found that, as for VBM in SPM, the results from SBM in FreeSurfer and did not replicate across the two datasets. Secondly, we conducted an ANOVAs with VBM in SPM combining both datasets, which represents the largest study to date of GMV in RD in participants matched on age, sex, SES and nonverbal reasoning (N=262) and controlling for these variables together with total GMV in the analysis. The main effect of Reading Ability revealed more GMV in the Control than the RD Group in left fusiform gyrus (medial and anterior to the visual word form area, VWFA), right superior temporal gyrus, right caudate, and left and right cerebellar lobule VI. It also revealed relatively less GMV in the Control Group in right middle orbitofrontal gyrus. However, there were no differences in left temporoparietal and inferior frontal regions (associated with spoken language) or in the visual word form area of the fusiform gyrus (associated with word recognition), inconsistent with the prevailing brain-based model of dyslexia (Pugh et al., 2000). We also found that the number of GMV differences between those with and without RD independent of dataset (main effect of Reading Ability) was dwarfed by those when comparing the two datasets independent of reading ability (main effect of Dataset). Lastly, the interaction effect identified four regions, including left inferior frontal gyrus, that showed GMV differences in RD that were dependent on the dataset, providing statistical confirmation of the lack of replication of these regions across the two datasets, despite the similarity of the groups’ overall profiles. Third, we investigated the effects of controlling for key variables, and, critically, found more GMV differences (including regions in language cortex) between the Control and RD Groups when total GMV was not controlled for. Omitting total GMV as a covariate of no interest also resulted in no interaction results, demonstrating lowered sensitivity for regions that did not replicate. This supports the notion that prior findings in the literature may in part have been a product of a suboptimal analysis approach frequently used in older studies (Eckert et al., 2016). Taken together, the results from this direct discovery and replication study followed by reproducibility studies demonstrate that the concerns raised by Ramus and colleagues (2018), about low reproducibility of GMV differences in RD, are valid.

### 4.1 Discovery and Replication Study

The between-group comparisons conducted for the Discovery Dataset and the Replication Dataset used VBM in SPM, consistent with most prior GMV studies of RD, and a widely used statistical threshold (p < 0.005) that is more conservative than those used in most of the 13 RD studies in the meta-analyses. This revealed several regions of more GMV in the Control compared to the RD Group. All of these regions have previously been reported to differ in RD, but none of these were born out of both datasets: left inferior temporal gyrus and cerebellum for the Discovery Dataset, and left inferior orbitofrontal, middle temporal, and lingual gyri and putamen for the Replication Dataset. There was more GMV in the Control Group in the right superior temporal gyrus in both datasets, but in different locations of the gyrus. The Discovery Dataset also revealed a region in the right middle orbitofrontal gyrus where the Control Group had relatively less GMV than the RD Group. When a more stringent threshold was applied to the analyses (p < 0.001), these results disappeared, with only one between-group difference observed in the Discovery Dataset for the left calcarine cortex.

### 4.2 Reproducibility Studies

Prior work has addressed the question whether the use of different analysis programs yield the same findings for between-group comparisons (Clausenius-Kalman, 2021, in studies of bilinguals versus monolinguals) and found the results to be affected by these varying computational frameworks. Our first Reproducibility Study tested for GMV differences between the Control and RD Groups using FreeSurfer (instead of VBM in SPM) and again found no replication of between-group differences in GMV across the Discovery and Replication Datasets. The analysis of GMV with FreeSurfer also revealed Controls to have more GMV than the RD Group in the left anterior transverse collateral sulcus in the Discovery Dataset, while there were no findings in the Replication Dataset. As such, the FreeSurfer analysis was similar to the VBM analysis in SPM with the conservative voxel-wise threshold (p < 0.001), both yielding few between-group difference in the Discovery Dataset, but not the same region. The results from this analysis demonstrate that (i) results (for greater GMV in Controls) did not replicate across the two datasets, and (ii) results from FreeSurfer show poor replication of those from VBM in SPM, which raises additional challenges for the field.

The second Reproducibility Study combined both datasets, resulting in a large (130 Control and 132 RD participants), well-controlled (for age, sex, SES, and tGMV) study. Results were generated using a statistical threshold of p < 0.005, which is more conservative than most studies in the meta-analyses (albeit less conservative than recommended cluster-forming thresholds for fMRI, see Woo et al., 2014) and well suited for reproducibility testing vis-à-vis entering covariates of no interest in the analysis. Here we discuss the results of the first ANOVA, and then those that followed to test for the role of covariates of no interest.

The ANOVA revealed that a main effect of Reading Ability (Discovery and Replication Datasets combined) on GMV, with more GMV in the Control than the RD Group in left fusiform gyrus (extending into parahippocampal gyrus and not including the VWFA), right superior temporal gyrus, and right caudate, as well as left and right cerebellar lobules VI. The main effect also revealed less GMV in the Control than the RD Groups in the right middle orbitofrontal gyrus. While there were not the expected findings in left orbitofrontal/inferior frontal and left temporoparietal cortices, other regions emerged from these results that have been reported in at least one of the above-noted meta-analyses on dyslexia, and each will be discussed in turn. The right superior temporal gyrus (STG) was in two meta-analyses (Linkersdorfer et al., 2012; Richlan et al., 2013), and while the location of the right STG in the present study is close to that reported by Richlan et al. (2012), it is noteworthy that another nearby section of the STG differed between the two datasets, and a third differed in RD depending on the dataset (i.e. the interaction, as discussed below), suggesting that the STG harbors variability, which in turn may give rise to spurious between-group differences. Our results in bilateral cerebellar lobule VI fit with the meta-analysis result by Linkersdorfer et al. (2012), but only partially with that by Eckert et al. (2016) who reported only right cerebellum, and not with that by Richlan et al. (2012), who did not identify differences in the cerebellum. While more GMV for the Control Group in left fusiform gyrus (FG) in the current study may appear to agree with the meta-analysis by Linkersdorfer et al. (2012), the location of the FG in the present study is more medial and anterior than that previously reported, thereby placing it outside of the VWFA (Cohen et al., 2000), and failing to support the theory that the VWFA is aberrant in RD. While the meta-analyses discussed above did not report on the caudate, a meta-analysis of GMV in RD which included logographic writing systems, did report left and right caudate differences (McGrath & Stoodley, 2019), and we found only the right caudate. In addition, our study revealed less GMV in the Control than the RD Group in right middle orbitofrontal gyrus, while prior meta-analyses generally only found regions of more GMV in controls (Linkersdorfer et al., 2012; Richlan et al., 2013; Eckert et al., 2016; however, see McGrath & Stoodley, 2019). Some of the original empirical studies that identified regions with more GMV in RD discussed these as playing a potential compensatory role.

Next, the main effect of Dataset revealed a substantial number of GMV differences between the two datasets despite these being matched on multiple demographic variables. This highlights the challenges of identifying GMV differences that are specific to RD among these more global discrepancies. It is notable that left IFG and left STG (extending into supramarginal and angular gyri) are among these, given their prevalent role in reading dyslexia and their purported impairment in dyslexia (Eckert et al., 2016; Linkersdorfer et al., 2012).

Lastly, the interaction between Reading Ability and Dataset on GMV differences in RD indicated four brain regions that emerged with more GMV in the Control Group for one dataset but not the other dataset: left inferior frontal, middle temporal, lingual and right superior temporal gyri. These were among the regions that were noted in either the Discovery or the Replication dataset (discussed above), but not both, and were hereby confirmed quantitatively as not being reproducible between these two datasets. While we discussed above the absence of left hemisphere language areas for those with RD despite the large sample size, left inferior frontal and middle temporal gyri were among these interaction results, suggesting that while they may differ in RD in one cohort, these differences do not manifest dependably in another cohort. This result provides an important lens by which to view the variation of results among past studies of dyslexia.

Next, we repeated this ANOVA three more times to understand how reproducible these results are when altering the covariates of no interest. We found that when we no longer controlled for age, sex, and SES, but solely controlled for total GMV, the results were similar to the original ANOVA, which may not be entirely be unexpected given that the Control and RD Groups were already equated on most variables, but nevertheless important given the relationships of these variables with GMV. Notably, this analysis (just like the original ANOVA) did not reveal differences in the RD Group in left cortex associated with spoken language, but the next two ANOVAs, which did not control for tGMV, but instead for tICV or for nothing at all, resulted in maps that included language regions (left temporal and frontal regions) within increasingly larger clusters. This suggests that prior studies may have identified left hemisphere language regions by not controlling for tGMV. Of the 13 meta-analysis studies, nine compared tGMV between the groups and of these six found no difference, while three found more tGMV in the control than the RD groups. Surprisingly, we found mean tGMV values to be higher in the Control Groups than the RD Groups for both datasets. While these results are contrary to prior studies, the main point is that there is a need to control for tGMV. For example, Eckert and colleagues reported on a retroactive study of 293 participants (164 RD and 129 Controls) drawn from multiple sites, and found less GMV in RD in left posterior superior temporal sulcus and orbitofrontal cortex using p < 0.001 uncorrected (Eckert et al., 2016). However, these findings disappeared when controlling for total GMV or FWE correction, underscoring the likelihood that many prior studies would have had no results if they had applied these more stringent analysis tactics. The current study, while similar in size, did identify multiple differences in RD despite controlling for total GMV and conducting FWE correction at a voxel-wise threshold of p < 0.005. but not at p < 0.001.

### 4.3 Are there GMV differences in RD?

The absence of robust, replicable differences in GMV left hemisphere cortical language regions raises the important question whether prior studies of GMV have mischaracterized the neuroanatomical bases of dyslexia. To answer this, it has to be considered that such cortical differences in GMV in left language cortex may well exist, but because they vary within and between participants in terms of their location and degree of severity, they cannot be detected reliably in group averages (beyond the GMV differences that exist in the general population). As such, they emerge for some studies and not others, even when participants in those studies have the same profiles, such as our Discovery and Replication Datasets. In a similar vein, the GMV differences between datasets (where groups without and with RD are combined) proved to be more numerous than one might expect, with a uniform distribution in terms of the direction of the effect (both less and more GMV for the Discovery versus Replication Datasets) and their locations (across all lobes and both hemispheres). These wide-ranging discrepancies in GMV amongst groups that were chosen under the same criteria, shed light on a major challenge in conducting GMV studies.

The ABCD Study is a multisite effort and such studies are susceptible to site-specific variations in participant recruitment, testing protocols and scanner characteristics, which can impact MRI data (Dias et al., 2022). However, unlike retrospective studies, the ABCD Study was designed prospectively and participant recruitment and testing adhered to the same protocol (Casey et al., 2019). Nevertheless, sites vary in their populations and equipment, and therefore our analyses included study site as a covariate of no interest. The main effects analysis results of GMV differences attributed to Dataset therefore reflect normal variation in population characteristics that exist despite a unified study protocol and selection of groups matched on demographics and behavioral characteristics. Participants were 9-10 years of age, raising the question as to whether these findings generalize to other age groups. Cortical morphology changes during childhood and adolescence (Sowell et al., 2003; Tamnes et al., 2017) and image quality is more variable in younger participants (Dias et al., 2022), primarily due to head motion (Blumenthal et al., 2002; Brown et al., 2010). The current study, therefore, offers a snapshot in time, and future studies could investigate the same sample when they are older. It is noteworthy that 9-10 years of age is well suited for the study of RD given that multiple prior studies reporting neuroanatomical differences in RD involved participants of this age (Jednoróg et al. 2014; Jednoróg et al. 2015; Krafnick et al., 2014; Ligges et al., 2022). Further, one study directly investigated GMV differences in RD at different ages, reporting findings for age 10 years, but not 13 years, nor for adults (Ligges et al., 2022). This suggests that the age of participants in the current study is optimal, coinciding with the age at which GMV differences in RD manifest robustly.

### 4.4 Implications and Future Directions

Taken together, the results from our direct Discovery and Replication Study as well as our Reproducibility Studies provide little support for less GMV in RD in left hemisphere language regions or elsewhere. Our Discovery and Replication Study indicates that poor replication is observed independent of the analysis program used to gauge GMV. Our Reproducibility Studies results indicated that earlier findings of GMV differences in RD may have come about (among other factors) by not controlling for global GMV. These findings validate the concerns expressed by Ramus and colleagues (2018) and indicate a need for considerable caution when considering results from prior GMV studies in dyslexia. For example, it would seem ill-advised to create a regions-of-interest for interrogation into brain function based on these older findings and the related meta-analyses. The widely accepted phenomenon of aberrant left-hemisphere language regions in dyslexia may still be correct, but it is not possible to capture these with GMV reliably, even when matching participant characteristics and optimizing analysis methods. Therefore, future studies need to include other measures, such as cortical thickness and sulcal depth to address this question with multiple, more informative metrics.

## 5 Conclusions

We conducted a direct Discovery and Replication Study of GMV differences in large groups of children with and without reading disability (dyslexia), using a more stringent methodological approach than those in past studies. We found no replication across the two datasets for differences in GMV between the Control and RD Groups. One Reproducibility Study repeated the Discovery and Replication Study this time using SBM in FreeSurfer instead of VBM in SPM, and also found a lack of replication of GMV differences between the two groups in both datasets. Another Reproducibility Study revealed that while there were no left language cortex differences even when the two datasets were combined, not controlling for total GMV resulted in more extensive clusters and, with it, language cortex. This may be one reason behind the reporting of these brain areas in prior GMV studies of RD. Based on these results, GMV differences in RD, if they exist, may not be captured with the metric of GMV, likely due to heterogeneous manifestation in RD as well as general variations in GMV among any two cohorts. As such, results for GMV differences in dyslexia cannot be considered to be reliable and the use of better neuroanatomical metrics in future studies is important.

## Acknowledgements

Data used in the preparation of this article were obtained from the Adolescent Brain Cognitive Development (ABCD) Study (https://abcdstudy.org), held in the NIMH Data Archive (NDA). This is a multisite, longitudinal study designed to recruit more than 10,000 children age 9–10 and follow them over 10 years into early adulthood. The ABCD Study® is supported by the National Institutes of Health and additional federal partners under award numbers U01DA041048, U01DA050989, U01DA051016, U01DA041022, U01DA0510 18, U01DA051037, U01DA050987, U01DA041174, U01DA041106, U01DA041117, U01DA041028, U01DA041134, U01DA050988, U01DA051039, U01DA041156, U01DA 041025, U01DA041120, U01DA051038, U01DA041148, U01DA041093, U01DA041089, U24DA041123, U24DA041147. A full list of supporters is available at https://abcdstudy.org/federal-partners.html.

## Data Availability Statement

All data files are available from the ABCD Study Data Repository from the NIMH Data Archive (https://nda.nih.gov/abcd). The DOI for this NDA Study is 10.15154/sr36-8f36.

## Ethics Approval Statement

The ABCD Study was performed under the principles of the Declaration of Helsinki (Garavan et al., 2018).

## Patient Consent Statement

Parent consent and child assent was obtained by the ABCD Study (Garavan et al., 2018).

## Permission to reproduce material from other sources

N/A

## Clinical Trial Registration

N/A

## References

Ashburner, J. (2007). A fast diffeomorphic image registration algorithm. NeuroImage, 38(1), 95–113. 10.1016/j.neuroimage.2007.07.007

Ashburner, J. (2015). VBM Tutorial. https://www.fil.ion.ucl.ac.uk/~john/misc/VBMclass15.pdf

Ashburner, J., & Friston, K. J. (2000). Voxel-Based Morphometry—The Methods. NeuroImage, 11(6), 805–821. 10.1006/nimg.2000.0582

Brambati, S. M., Termine, C., Ruffino, M., Stella, G., Fazio, F., Cappa, S. F., & Perani, D. (2004). Regional reductions of gray matter volume in familial dyslexia. Neurology, 63(4), 742–745. 10.1212/01.WNL.0000134673.95020.EE

Brito, N. H., & Noble, K. G. (2018). The independent and interacting effects of socioeconomic status and dual-language use on brain structure and cognition. Developmental Science, 21(6), e12688–e12688. PubMed. 10.1111/desc.12688

Blumenthal, J. D., Zijdenbos, A., Molloy, E., & Giedd, J. N. (2002). Motion artifact in magnetic resonance imaging: Implications for automated analysis. NeuroImage, 16(1), 89–92. 10.1006/nimg.2002.1076

Brown, T. T., Kuperman, J. M., Erhart, M., White, N. S., Roddey, J. C., Shankaranarayanan, A., Han, E. T., Rettmann, D., & Dale, A. M. (2010). Prospective motion correction of high-resolution magnetic resonance imaging data in children. NeuroImage, 53(1), 139–145. 10.1016/j.neuroimage.2010.06.017

Brown, W. E., Eliez, S., Menon, V., Rumsey, J. M., White, C. D., & Reiss, A. L. (2001). Preliminary evidence of widespread morphological variations of the brain in dyslexia. Neurology, 56(6), 781–783. 10.1212/WNL.56.6.781

Casey BJ, Cannonier T, Conley MI, Cohen AO, Barch DM, Heitzeg MM, Soules ME, Teslovich T, Dellarco DV, Garavan H, Orr CA, Wager TD, Banich MT, Speer NK, Sutherland MT, Riedel MC, Dick AS, Bjork JM, Thomas KM, Chaarani B, Mejia MH, Hagler DJ Jr, Daniela Cornejo M, Sicat CS, Harms MP, Dosenbach NUF, Rosenberg M, Earl E, Bartsch H, Watts R, Polimeni JR, Kuperman JM, Fair DA, Dale AM; ABCD Imaging Acquisition Workgroup. The Adolescent Brain Cognitive Development (ABCD) study: Imaging acquisition across 21 sites. Dev Cogn Neurosci. 2018 Aug;32:43–54. doi: 10.1016/j.dcn.2018.03.001. Epub 2018 Mar 14. PMID: 29567376; PMCID: PMC5999559.

Catts, H. W., Adlof, S. M., Hogan, T. P., & Weismer, S. E. (2005). Are Specific Language Impairment and Dyslexia Distinct Disorders? *Journal of Speech*, Language, and Hearing Research, 48(6), 1378–1396. 10.1044/1092-4388(2005/096)

Claussenius Kalman, H., Vaughn, K. A., Archila Suerte, P., & Hernandez, A. E. (2020). Age of acquisition impacts the brain differently depending on neuroanatomical metric. Human Brain Mapping, 41(2), 484–502.

Cohen L, Dehaene S, Naccache L, Lehéricy S, Dehaene-Lambertz G, Hénaff MA, Michel F. The visual word form area: spatial and temporal characterization of an initial stage of reading in normal subjects and posterior split-brain patients. Brain. 2000 Feb;123 (Pt 2):291–307. doi: 10.1093/brain/123.2.291. PMID: 10648437.

Drake, W. E. (1968). Clinical and Pathological Findings in a Child with a Developmental Learning Disability. Journal of Learning Disabilities, 1(9), 486–502. 10.1177/002221946800100901

Dias MFM, Carvalho P, Castelo-Branco M, & Valente Duarte J. Cortical thickness in brain imaging studies using FreeSurfer and CAT12: A matter of reproducibility. Neuroimage Rep. 2022 Oct 7;2(4):100137. doi: 10.1016/j.ynirp.2022.100137. PMID: 40567565; PMCID: PMC12172744.

Eckert, M. A., Berninger, V. W., Vaden, K. I., Gebregziabher, M., & Tsu, L. (2016). Gray Matter Features of Reading Disability: A Combined Meta-Analytic and Direct Analysis Approach(1,2,3,4). eNeuro, 3(1). 10.1523/ENEURO.0103-15.2015

Eckert, M. A., Leonard, C. M., Wilke, M., Eckert, M., Richards, T., Richards, A., & Berninger, V. (2005). Anatomical Signatures of Dyslexia in Children: Unique Information from Manual and Voxel Based Morphometry Brain Measures. Cortex, 41(3), 304–315. 10.1016/S0010-9452(08)70268-5

Evans, T. M., Flowers, D. L., Napoliello, E. M., & Eden, G. F. (2014). Sex-specific Gray Matter Volume Differences in Females with Developmental Dyslexia. Brain Structure & Function, 219(3), 1041–1054. 10.1007/s00429-013-0552-4

Fürtjes, A. E., Cole, J. H., Couvy-Duchesne, B., & Ritchie, S. J. (2023). A quantified comparison of cortical atlases on the basis of trait morphometricity. Cortex, 158, 110–126. 10.1016/j.cortex.2022.11.001

Garavan, H., Bartsch, H., Conway, K., Decastro, A., Goldstein, R. Z., Heeringa, S., Jernigan, T., Potter, A., Thompson, W., & Zahs, D. (2018). Recruiting the ABCD sample: Design considerations and procedures. Developmental Cognitive Neuroscience, 32, 16–22. 10.1016/j.dcn.2018.04.004

Germanò, E., Gagliano, A., & Curatolo, P. (2010). Comorbidity of ADHD and Dyslexia. Developmental Neuropsychology, 35(5), 475–493. 10.1080/87565641.2010.494748

Gershon, R. C., Cook, K. F., Mungas, D., Manly, J. J., Slotkin, J., Beaumont, J. L., & Weintraub, S. (2014). Language Measures of the NIH Toolbox Cognition Battery. Journal of the International Neuropsychological Society, 20(6), 642–651. 10.1017/S1355617714000411

Gershon, R. C., Slotkin, J., Manly, J. J., Blitz, D. L., Beaumont, J. L., Schnipke, D., Wallner-Allen, K., Golinkoff, R. M., Gleason, J. B., Hirsh-Pasek, K., Adams, M. J., & Weintraub, S. (2013). IV. NIH TOOLBOX COGNITION BATTERY (CB): MEASURING LANGUAGE (VOCABULARY COMPREHENSION AND READING DECODING): NIH TOOLBOX COGNITION BATTERY (CB). Monographs of the Society for Research in Child Development, 78(4), 49–69. 10.1111/mono.12034

Hoeft, F., Meyler, A., Hernandez, A., Juel, C., Taylor-Hill, H., Martindale, J. L., McMillon, G., Kolchugina, G., Black, J. M., Faizi, A., Deutsch, G. K., Siok, W. T., Reiss, A. L., Whitfield-Gabrieli, S., & Gabrieli, J. D. E. (2007). Functional and morphometric brain dissociation between dyslexia and reading ability. Proceedings of the National Academy of Sciences of the United States of America, 104(10), 4234–4239. 10.1073/pnas.0609399104

Hoeft, F., Ueno, T., Reiss, A. L., Meyler, A., Whitfield-Gabrieli, S., Glover, G. H., Keller, T. A., Kobayashi, N., Mazaika, P., Jo, B., Just, M. A., & Gabrieli, J. D. E. (2007). Prediction of children’s reading skills using behavioral, functional, and structural neuroimaging measures. Behavioral Neuroscience, 121(3), 602–613. 10.1037/0735-7044.121.3.602

Iqbal, S. A., Wallach, J. D., Khoury, M. J., Schully, S. D., & Ioannidis, J. P. A. (2016). Reproducible Research Practices and Transparency across the Biomedical Literature. PLOS Biology, 14(1), e1002333. 10.1371/journal.pbio.1002333

Jednoróg, K., Gawron, N., Marchewka, A., Heim, S., & Grabowska, A. (2014). Cognitive subtypes of dyslexia are characterized by distinct patterns of grey matter volume. Brain Structure and Function, 219(5), 1697–1707. 10.1007/s00429-013-0595-6

Jednoróg K, Marchewka A, Altarelli I, Monzalvo Lopez AK, van Ermingen-Marbach M, Grande M, Grabowska A, Heim S, Ramus F. How reliable are gray matter disruptions in specific reading disability across multiple countries and languages? Insights from a large-scale voxel-based morphometry study. Hum Brain Mapp. 2015 May;36(5):1741–54. doi: 10.1002/hbm.22734. Epub 2015 Jan 17. PMID: 25598483; PMCID: PMC6869714.

Karcher, N. R., & Barch, D. M. (2021). The ABCD study: Understanding the development of risk for mental and physical health outcomes. Neuropsychopharmacology, 46(1), 131–142. 10.1038/s41386-020-0736-6

Katusic, S. K., Colligan, R. C., Barbaresi, W. J., Schaid, D. J., & Jacobsen, S. J. (2001). Incidence of Reading Disability in a Population-Based Birth Cohort, 1976–1982, Rochester, Minn. Mayo Clinic Proceedings, 76(11), 1081–1092. 10.4065/76.11.1081

Kline, A., & Luo, Y. (2022). PsmPy: A Package for Retrospective Cohort Matching in Python [Computer software]. https://pypi.org/project/psmpy/

Krafnick, A. J., Flowers, D. L., Luetje, M. M., Napoliello, E. M., & Eden, G. F. (2014). An investigation into the origin of anatomical differences in dyslexia. The Journal of Neuroscience: The Official Journal of the Society for Neuroscience, 34(3), 901–908. 10.1523/JNEUROSCI.2092-13.2013

Kronbichler, M., Wimmer, H., Staffen, W., Hutzler, F., Mair, A., & Ladurner, G. (2008). Developmental dyslexia: Gray matter abnormalities in the occipitotemporal cortex. Human Brain Mapping, 29(5), 613–625. 10.1002/hbm.20425

Liloia, D., Crocetta, A., Cauda, F., Duca, S., Costa, T., & Manuello, J. (2022). Seeking Overlapping Neuroanatomical Alterations between Dyslexia and Attention-Deficit/Hyperactivity Disorder: A Meta-Analytic Replication Study. Brain Sciences, 12(10), 1367. 10.3390/brainsci12101367

Ligges C, Ligges M, & Gaser C. Cross-Sectional Investigation of Brain Volume in Dyslexia. Front Neurol. 2022 Mar 8;13:847919. doi: 10.3389/fneur.2022.847919. PMID: 35350399; PMCID: PMC8957969.

Linkersdörfer, J., Lonnemann, J., Lindberg, S., Hasselhorn, M., & Fiebach, C. J. (2012). Grey Matter Alterations Co-Localize with Functional Abnormalities in Developmental Dyslexia: An ALE Meta-Analysis. PLOS ONE, 7(8), e43122. 10.1371/journal.pone.0043122

Liu, L., You, W., Wang, W., Guo, X., Peng, D., & Booth, J. (2013). Altered brain structure in Chinese dyslexic children. Neuropsychologia, 51(7), 1169–1176. 10.1016/j.neuropsychologia.2013.03.010

Lotze M, Domin M, Schmidt CO, Hosten N, Grabe HJ, Neumann N. Income is associated with hippocampal/amygdala and education with cingulate cortex grey matter volume. Sci Rep. 2020 Nov 2;10(1):18786. doi: 10.1038/s41598-020-75809-9. PMID: 33139786; PMCID: PMC7608615.

McDaniel, M. (2005). Big-brained people are smarter: A meta-analysis of the relationship between in vivo brain volume and intelligence. Intelligence, 33(4), 337–346. 10.1016/j.intell.2004.11.005

McGrath, L. M., & Stoodley, C. J. (2019). Are there shared neural correlates between dyslexia and ADHD? A meta-analysis of voxel-based morphometry studies. Journal of Neurodevelopmental Disorders, 11(1), 31. 10.1186/s11689-019-9287-8

Menghini, D., Hagberg, G. E., Petrosini, L., Bozzali, M., Macaluso, E., Caltagirone, C., & Vicari, S. (2008). Structural correlates of implicit learning deficits in subjects with developmental dyslexia. Annals of the New York Academy of Sciences, 1145, 212–221. 10.1196/annals.1416.010

Pernet, C. R., Poline, J. B., Demonet, J. F., & Rousselet, G. A. (2009). Brain classification reveals the right cerebellum as the best biomarker of dyslexia. BMC Neuroscience, 10, 67. 10.1186/1471-2202-10-67

Pugh, K. R., Mencl, W. E., Jenner, A. R., Katz, L., Frost, S. J., Lee, J. R., et al. (2001). Neurobiological studies of reading and reading disability. Journal of Communication Disorders, 34(6), 479–492. doi.org/10.1016/S0021-9924(01)00060-0

Radua, J., Canales-Rodríguez, E. J., Pomarol-Clotet, E., & Salvador, R. (2014). Validity of modulation and optimal settings for advanced voxel-based morphometry. NeuroImage, 86, 81–90. 10.1016/j.neuroimage.2013.07.084

Rakesh D, Whittle S. Socioeconomic status and the developing brain - A systematic review of neuroimaging findings in youth. Neurosci Biobehav Rev. 2021 Nov;130:379–407. doi: 10.1016/j.neubiorev.2021.08.027. Epub 2021 Aug 30. PMID: 34474050.

Ramus, F., Altarelli, I., Jednoróg, K., Zhao, J., & Scotto di Covella, L. (2018). Neuroanatomy of developmental dyslexia: Pitfalls and promise. Neuroscience & Biobehavioral Reviews, 84, 434–452. 10.1016/j.neubiorev.2017.08.001

Raschle, N. M., Chang, M., & Gaab, N. (2011). Structural brain alterations associated with dyslexia predate reading onset. NeuroImage, 57(3), 742–749. 10.1016/j.neuroimage.2010.09.055

Richlan, F. (2014). Functional neuroanatomy of developmental dyslexia: The role of orthographic depth. Frontiers in Human Neuroscience, 8. 10.3389/fnhum.2014.00347

Richlan, F., Kronbichler, M., & Wimmer, H. (2012). Structural abnormalities in the dyslexic brain: A meta-analysis of voxel-based morphometry studies. Human Brain Mapping, 34(11), 3055–3065. 10.1002/hbm.22127

Sandak, R., Mencl, W. E., Frost, S. J., and Pugh, K. R. (2004). The neurobiological basis of skilled and impaired reading: Recent findings and new directions. Scientific Studies of Reading, 8, 273–292. DOI: 10.1207/S1532799Xssr0803_6

Shaywitz, S. E. (1998). Dyslexia. New England Journal of Medicine, 338(5), 307–312. 10.1056/NEJM199801293380507

Silani, G., Frith, U., Demonet, J.-F., Fazio, F., Perani, D., Price, C., Frith, C. D., & Paulesu, E. (2005). Brain abnormalities underlying altered activation in dyslexia: A voxel based morphometry study. Brain, 128(10), 2453–2461. 10.1093/brain/awh579

Siok, W. T., Niu, Z., Jin, Z., Perfetti, C. A., & Tan, L. H. (2008). A structural–functional basis for dyslexia in the cortex of Chinese readers. Proceedings of the National Academy of Sciences, 105(14), 5561–5566. 10.1073/pnas.0801750105

Sowell, E. R., Peterson, B. S., Thompson, P. M., Welcome, S. E., Henkenius, A. L., & Toga, A. W. (2003). Mapping cortical change across the human life span. Nature Neuroscience, 6(3), 309–315. 10.1038/nn1008

Tamnes, C. K., Herting, M. M., Goddings, A.-L., Meuwese, R., Blakemore, S.-J., Dahl, R. E., Güroğlu, B., Raznahan, A., Sowell, E. R., Crone, E. A., & Mills, K. L. (2017). Development of the Cerebral Cortex across Adolescence: A Multisample Study of Inter-Related Longitudinal Changes in Cortical Volume, Surface Area, and Thickness. The Journal of Neuroscience, 37(12), 3402–3412. 10.1523/JNEUROSCI.3302-16.2017

Steinbrink, C., Vogt, K., Kastrup, A., Müller, H.-P., Juengling, F. D., Kassubek, J., & Riecker, A. (2008). The contribution of white and gray matter differences to developmental dyslexia: Insights from DTI and VBM at 3.0T. Neuropsychologia, 46(13), 3170–3178. 10.1016/j.neuropsychologia.2008.07.015

Tamboer, P., Scholte, H. S., & Vorst, H. C. M. (2015). Dyslexia and voxel-based morphometry: Correlations between five behavioural measures of dyslexia and gray and white matter volumes. Annals of Dyslexia, 65(3), 121–141. 10.1007/s11881-015-0102-2

Vinckenbosch, E., Robichon, F., & Eliez, S. (2005). Gray matter alteration in dyslexia: Converging evidence from volumetric and voxel-by-voxel MRI analyses. Neuropsychologia, 43(3), 324–331. 10.1016/j.neuropsychologia.2004.06.023

Wilkinson, G. S., & Robertson, G. J. (2006). *WRAT 4: Wide range achievement test*. Psychological Assessment Resources Lutz, FL.

Woo CW, Krishnan A, Wager TD. Cluster-extent based thresholding in fMRI analyses: pitfalls and recommendations. Neuroimage. 2014 May 1;91:412–9. doi: 10.1016/j.neuroimage.2013.12.058. Epub 2014 Jan 8. PMID: 24412399; PMCID: PMC4214144.

Xia, Z., Hoeft, F., Zhang, L., & Shu, H. (2016). Neuroanatomical anomalies of dyslexia: Disambiguating the effects of disorder, performance, and maturation. Neuropsychologia, 81, 68–78. 10.1016/j.neuropsychologia.2015.12.003

Yang, Y.-H., Yang, Y., Chen, B.-G., Zhang, Y.-W., & Bi, H.-Y. (2016). Anomalous Cerebellar Anatomy in Chinese Children with Dyslexia. Frontiers in Psychology, 7. 10.3389/fpsyg.2016.00324

