## Supplemental Information for "A discovery and replication study of dyslexia does not reveal reproducible gray matter volume differences"

Supplement

**Supplementary Table 1. ANOVA (VBM): Main Effect of Reading Ability, Main Effect of Dataset, and their Interactions with Total GMV as a Covariate**

| **Main Effect of Reading Ability** | | | | | | |
| --- | --- | --- | --- | --- | --- | --- |
| **x** | **y** | **z** | **Anatomical Region** | **BA** | **Z** | **kE** |
| **Control > RD** | | | | | | |
| -12 | -64 | -24 | Left Cerebellum Lobule VI |  | 3.83 | 1036 |
| -16 | -58 | -32 | Left Cerebellum Lobule VI |  | 3.69 |  |
| -22 | -72 | -24 | Left Cerebellum Lobule VI |  | 2.69 |  |
| 15 | -2 | 26 | Right Caudate | 48 | 3.4 | 297 |
| 14 | 15 | 20 | Right Caudate | 48 | 3.25 |  |
| 21 | 6 | 16 | Right Caudate | 48 | 3.01 |  |
| 52 | -36 | 12 | Right Superior Temporal Gyrus | 22 | 3.57 | 354 |
| 56 | -45 | 9 | Right Middle Temporal Gyrus | 22 | 3.29 |  |
| 45 | -44 | 4 | Right Middle Temporal Gyrus | 21 | 2.85 |  |
| 12 | -62 | -21 | Right Cerebellum Lobule VI |  | 3.05 | 291 |
| **Control < RD** | | | | | | |
|  |  |  | None |  |  |  |
| **Main Effect of Dataset** | | | | | | |
| **x** | **y** | **z** | **Anatomical Region** | **BA** | **Z** | **kE** |
| **Discovery > Replication** | | | | | | |
| -14 | 18 | 68 | Left Supplementary Motor Area | 6 | 4.39 | 1663 |
| 0 | -3 | 70 | Left Supplementary Motor Area | 6 | 4.27 |  |
| 3 | 20 | 64 | Right Supplementary Motor Area | 6 | 3.55 |  |
| -69 | -15 | -21 | Left Middle Temporal Gyrus | 21 | 7.8 | 11333 |
| -68 | -22 | -27 | Left Inferior Temporal Gyrus | 20 | 7.37 |  |
| -70 | -20 | -14 | Left Middle Temporal Gyrus | 21 | 7.09 |  |
| -33 | -78 | -56 | Left Cerebellum Lobule VIIb | 19 | 4.23 | 456 |
| -20 | -76 | -58 | Left Cerebellum Lobule VIIb | 18 | 4.07 |  |
| -40 | -70 | -57 | Left Cerebellum Lobule VIIb | 37 | 4.02 |  |
| -4 | -98 | -16 | Left Lingual Gyrus | 18 | 4.6 | 3478 |
| -45 | -86 | -15 | Left Inferior Occipital Gyrus | 19 | 4.44 |  |
| -3 | -100 | -9 | Left Calcarine Fissure | 18 | 4.41 |  |
| 63 | 4 | -20 | Right Middle Temporal Gyrus | 38 | Inf | 12013 |
| 63 | 2 | -28 | Right Middle Temporal Gyrus | 21 | 7.24 |  |
| 54 | 18 | -24 | Right Middle Temporal Pole | 38 | 7.02 |  |
| 14 | -12 | 60 | Right Supplementary Motor Area | 6 | 3.75 | 256 |
| 26 | -2 | 70 | Right Superior Frontal Gyrus | 6 | 2.97 |  |
| 9 | -27 | -6 | Right Lingual Gyrus | 50 | 4.14 | 366 |
| 8 | -34 | -3 | Right Lingual Gyrus | 36 | 3.91 |  |
| 3 | -27 | 6 | Right Thalamus | 50 | 3 |  |
| 3 | -63 | 12 | Right Calcarine Fissure | 18 | 3.61 | 428 |
| 6 | -66 | -57 | Right Cerebellum Lobule VIII |  | 5.12 | 596 |
| 0 | -58 | -60 | Right Cerebellum Lobule IX |  | 4.44 |  |
| 8 | -58 | -63 | Right Cerebellum Lobule IX |  | 4.34 |  |
| 28 | -75 | -60 | Right Cerebellum Lobule VIII |  | 4 | 475 |
| 28 | -81 | -54 | Right Cerebellum Lobule VIIb |  | 3.87 |  |
| 40 | -69 | -58 | Right Cerebellum Lobule VIIb |  | 3.75 |  |
| **Discovery < Replication** | | | | | | |
| -28 | -27 | 69 | Left Postcentral Gyrus | 4 | 4.67 | 1195 |
| -12 | -30 | 78 | Left Paracentral Lobule | 4 | 4.12 |  |
| -20 | -33 | 76 | Left Postcentral Gyrus | 1 | 3.8 |  |
| -18 | -34 | -45 | Left Cerebellum Lobule X |  | 5.23 | 543 |
| -12 | -38 | -50 | Left Cerebellum Lobule IX |  | 4.24 |  |
| -30 | -46 | -50 | Left Cerebellum Lobule VIII |  | 3.05 |  |
| -36 | -36 | -8 | Left Hippocampus | 54 | 4.26 | 786 |
| -30 | -45 | 0 | Left Lingual Gyrus | 19 | 3.96 |  |
| -21 | -34 | 8 | Left Hippocampus | 50 | 3.96 |  |
| -22 | -64 | -38 | Left Cerebellum Lobule VIII |  | 3.51 | 482 |
| -14 | -66 | -33 | Left Cerebellum Lobule VIII |  | 3.15 |  |
| -27 | -57 | -38 | Left Cerebellum Lobule VIII |  | 2.92 |  |
| -30 | -87 | 38 | Left Middle Occipital Gyrus | 19 | 5.6 | 8485 |
| -63 | -51 | 39 | Left Supramarginal Gyrus | 39 | 5.55 |  |
| -68 | -48 | 20 | Left Superior Temporal Gyrus | 39 | 5.28 |  |
| 4 | 28 | -32 | Right Gyrus Rectus | 11 | 5.21 | 946 |
| 4 | 56 | -28 | Right Gyrus Rectus | 11 | 4.44 |  |
| 6 | 51 | -20 | Right Gyrus Rectus | 11 | 4.05 |  |
| 33 | -14 | -2 | Right Putamen | 49 | 5.52 | 7093 |
| 0 | 6 | 4 | Left Caudate | 48 | 4.81 |  |
| -32 | -15 | 2 | Left Putamen | 49 | 4.61 |  |
| 70 | -26 | 14 | Right Superior Temporal Gyrus | 22 | 4.06 | 290 |
| 66 | -22 | 8 | Right Superior Temporal Gyrus | 41 | 3.42 |  |
| 21 | -34 | -44 | Right Cerebellum Lobule X |  | 4.78 | 1455 |
| 15 | -38 | -51 | Right Cerebellum Lobule IX |  | 4.04 |  |
| 28 | -51 | -52 | Right Cerebellum Lobule VIII |  | 3.72 |  |
| 32 | -44 | -2 | Right Fusiform Gyrus | 36 | 4.58 | 1175 |
| 38 | -38 | -8 | Right Parahippocampal Gyrus | 36 | 4.29 |  |
| 38 | -45 | -10 | Right Fusiform Gyrus | 37 | 3.81 |  |
| 12 | -46 | -27 | Right Cerebellum Lobule IV, V |  | 3.78 | 590 |
| -8 | -48 | -26 | Vermis X |  | 3.64 |  |
| 9 | -54 | -26 | Vermis IX |  | 3.56 |  |
| 34 | -84 | 33 | Right Middle Occipital Gyrus | 39 | 5.97 | 9386 |
| 30 | -90 | 15 | Right Middle Occipital Gyrus | 18 | 5.9 |  |
| 33 | -78 | 45 | Right Superior Occipital Gyrus | 7 | 5.76 |  |
| **Interaction of Reading Ability by Dataset** | | | | | | |
| **x** | **y** | **z** | **Anatomical Region** | **BA** | **Z** | **kE** |
| -38 | 34 | -24 | Left Inferior Orbitofrontal Gyrus | 47 | 3.67 | 345 |
| -52 | 42 | -10 | Left Inferior Orbitofrontal Gyrus | 47 | 3.6 |  |
| -40 | 30 | -10 | Left Inferior Orbitofrontal Gyrus | 47 | 3.19 |  |
| -45 | 10 | 8 | Left Inferior Frontal Gyrus pars Opercularis | 44 | 3.52 | 271 |
| -52 | 20 | -3 | Left Inferior Orbitofrontal Gyrus | 45 | 2.81 |  |
| -48 | -21 | -12 | Left Middle Temporal Gyrus | 21 | 4.15 | 707 |
| -45 | -28 | -10 | Left Middle Temporal Gyrus | 21 | 3.25 |  |
| -40 | -27 | -3 | Left Hippocampus | 48 | 3.23 |  |
| 36 | 32 | -24 | Right Inferior Orbitofrontal Gyrus | 47 | 3.59 | 597 |
| 22 | 16 | -21 | Right Inferior Orbitofrontal Gyrus | 47 | 3.39 |  |
| 22 | 36 | -24 | Right Superior Orbitofrontal Gyrus | 11 | 2.64 |  |
| 44 | -40 | 21 | Right Superior Temporal Gyrus | 22 | 3.7 | 331 |

*Note: Black font indicates peak coordinates of cluster and its cluster size, and gray font indicates sub-peak coordinates within that cluster. Coordinates indicating anatomical location are in Montreal Neurological Institute (MNI) stereotaxic space. See Figure 2*.

**Supplementary Table 2. ANOVA (VBM): Main Effect of Reading Ability, Main Effect of Dataset, and their Interactions with Total ICV as a Covariate**

| **Main Effect of Reading Ability** | | | | | | |
| --- | --- | --- | --- | --- | --- | --- |
| **x** | **y** | **z** | **Anatomical Region** | **BA** | **Z** | **kE** |
| **Control > RD** | | | | | | |
| -28 | 2 | 51 | Left Middle Frontal Gyrus | 6 | 3.55 | 451 |
| -20 | -6 | 48 | Left Superior Frontal Gyrus | 6 | 3.29 |  |
| -32 | -6 | 57 | Left Precentral Gyrus | 6 | 3.15 |  |
| -28 | -24 | 48 | Left Precentral Gyrus | 4 | 3.38 | 671 |
| -42 | -18 | 52 | Left Postcentral Gyrus | 4 | 3.04 |  |
| -16 | -22 | 60 | Left Precentral Gyrus | 6 | 3.02 |  |
| -70 | -26 | -3 | Left Middle Temporal Gyrus | 21 | 3.88 | 1028 |
| -62 | -22 | -2 | Left Middle Temporal Gyrus | 22 | 3.29 |  |
| -58 | -50 | 4 | Left Middle Temporal Gyrus | 21 | 3.2 |  |
| -12 | -64 | -24 | Left Cerebellum Lobule VI |  | 4.92 | 89409 |
| 14 | -54 | -28 | Right Cerebellum Lobule VI |  | 4.9 |  |
| -16 | -58 | -32 | Left Cerebellum Lobule VI |  | 4.71 |  |
| 30 | -14 | 48 | Right Precentral Gyrus | 6 | 3.84 | 717 |
| 30 | -22 | 46 | Right Postcentral Gyrus | 4 | 3.33 |  |
| 34 | -14 | 56 | Right Precentral Gyrus | 6 | 3.19 |  |
| 34 | -84 | 9 | Right Middle Occipital Gyrus | 19 | 3.53 | 595 |
| 30 | -78 | 2 | Right Fusiform Gyrus | 18 | 3.2 |  |
| 44 | -88 | -4 | Right Inferior Occipital Gyrus | 18 | 2.94 |  |
| **Control < RD** | | | | | | |
|  |  |  | None |  |  |  |
| **Main Effect of Dataset** | | | | | | |
| **x** | **y** | **z** | **Anatomical Region** | **BA** | **Z** | **kE** |
| **Discovery > Replication** | | | | | | |
| -27 | 36 | -24 | Left Middle Orbitofrontal Gyrus | 11 | 4.7 | 311 |
| -22 | 26 | -27 | Left Inferior Orbitofrontal Gyrus | 47 | 4.25 |  |
| -20 | 36 | -27 | Left Superior Orbitofrontal Gyrus | 11 | 3.9 |  |
| -69 | -15 | -21 | Left Middle Temporal Gyrus | 21 | 6.26 | 2883 |
| -68 | -22 | -27 | Left Inferior Temporal Gyrus | 20 | 6.05 |  |
| -66 | -8 | -27 | Left Inferior Temporal Gyrus | 21 | 5.58 |  |
| -8 | -100 | -14 | Left Calcarine Fissure | 18 | 3.68 | 255 |
| -3 | -102 | -4 | Left Calcarine Fissure | 18 | 3.3 |  |
| -15 | -99 | -18 | Left Lingual Gyrus | 18 | 3.08 |  |
| 21 | 32 | -27 | Right Superior Orbitofrontal Gyrus | 11 | 5 | 528 |
| 27 | 40 | -22 | Right Middle Orbitofrontal Gyrus | 11 | 4.29 |  |
| 28 | 28 | -26 | Right Inferior Orbitofrontal Gyrus | 47 | 4.23 |  |
| 63 | 4 | -22 | Right Middle Temporal Gyrus | 38 | 6.53 | 3578 |
| 68 | -9 | -26 | Right Middle Temporal Gyrus | 21 | 5.5 |  |
| 54 | 18 | -24 | Right Middle Temporal Pole | 38 | 5.48 |  |
| **Discovery < Replication** | | | | | | |
| -44 | 27 | 45 | Left Middle Frontal Gyrus | 8 | 4.57 | 1401 |
| -51 | 18 | 40 | Left Middle Frontal Gyrus | 8 | 4.16 |  |
| -32 | 39 | 45 | Left Middle Frontal Gyrus | 9 | 3.72 |  |
| -68 | -16 | 16 | Left Postcentral Gyrus | 1 | 3.82 | 474 |
| -52 | -9 | 20 | Left Postcentral Gyrus | 4 | 3.33 |  |
| -69 | -14 | 24 | Left Postcentral Gyrus | 1 | 2.93 |  |
| -26 | -28 | 70 | Left Postcentral Gyrus | 4 | 4.02 | 1050 |
| -20 | -33 | 76 | Left Postcentral Gyrus | 1 | 3.44 |  |
| -28 | -39 | 70 | Left Postcentral Gyrus | 5 | 3.43 |  |
| -8 | -28 | 27 | Left Posterior Cingulum | 23 | 3.95 | 997 |
| 4 | -9 | 26 | Right Middle Cingulum | 24 | 3.65 |  |
| -6 | -22 | 33 | Left Middle Cingulum | 23 | 3.52 |  |
| -18 | -34 | -45 | Left Cerebellum Lobule X |  | 4.85 | 6827 |
| -27 | -64 | -38 | Left Cerebellum Crus I |  | 4.45 |  |
| -12 | -38 | -50 | Left Cerebellum Lobule IX |  | 4.41 |  |
| 8 | 57 | -27 | Right Superior Orbitofrontal Gyrus | 11 | 4.87 | 6556 |
| 4 | 28 | -32 | Right Gyrus Rectus | 11 | 4.86 |  |
| 4 | 63 | -21 | Right Superior Orbitofrontal Gyrus | 11 | 4.83 |  |
| 12 | 36 | 34 | Right Middle Cingulum | 8 | 3.66 | 455 |
| 12 | 21 | 45 | Right Middle Cingulum | 8 | 3.37 |  |
| 14 | 14 | 50 | Right Superior Frontal Gyrus | 6 | 3.26 |  |
| 40 | -76 | 45 | Right Angular Gyrus | 39 | 6.79 | 68849 |
| 42 | -81 | 36 | Right Middle Occipital Gyrus | 39 | 6.49 |  |
| 50 | -74 | 38 | Right Angular Gyrus | 39 | 6.37 |  |
| **Interaction of Reading Ability by Dataset** | | | | | | |
| **x** | **y** | **z** | **Anatomical Region** | **BA** | **Z** | **kE** |
| -42 | -42 | 16 | Left Superior Temporal Gyrus | 22 | 3.77 | 320 |
| -38 | -38 | 24 | Left Rolandic Opercularis | 40 | 3.64 |  |
| -44 | -30 | 28 | Left Supramarginal Gyrus | 40 | 2.83 |  |
| 44 | -40 | 21 | Right Superior Temporal Gyrus | 22 | 3.81 | 484 |
| 40 | -45 | 28 | Right Angular Gyrus | 39 | 2.76 |  |
| 44 | -39 | 12 | Right Superior Temporal Gyrus | 22 | 2.65 |  |

*Note: Black font indicates peak coordinates of cluster and its cluster size, and gray font indicates sub-peak coordinates within that cluster. Coordinates indicating anatomical location are in Montreal Neurological Institute (MNI) stereotaxic space. See Figure 2*.

**Supplementary Table 3. ANOVA (VBM): Main Effect of Reading Ability, Main Effect of Dataset, and their Interactions with no Covariates**

| **Main Effect of Reading Ability** | | | | | | |
| --- | --- | --- | --- | --- | --- | --- |
| **x** | **y** | **z** | **Anatomical Region** | **BA** | **Z** | **kE** |
| **Control > RD** | | | | | | |
| 30 | -14 | 48 | Right Precentral Gyrus | 6 | 4.19 | 1634 |
| 22 | -12 | 51 | Right Precentral Gyrus | 6 | 3.73 |  |
| 36 | -21 | 50 | Right Postcentral Gyrus | 4 | 3.6 |  |
| 14 | -54 | -28 | Right Cerebellum Lobule VI |  | 5.16 | 2E+05 |
| -12 | -64 | -24 | Left Cerebellum Lobule VI |  | 5.13 |  |
| -16 | -58 | -32 | Left Cerebellum Lobule VI |  | 4.97 |  |
| **Control < RD** | | | | | | |
|  |  |  | None |  |  |  |
| **Main Effect of Dataset** | | | | | | |
| **x** | **y** | **z** | **Anatomical Region** | **BA** | **Z** | **kE** |
| **Discovery > Replication** | | | | | | |
| 0 | -3 | 70 | Left Supplementary Motor Area | 6 | 4.08 | 1192 |
| -14 | 18 | 68 | Left Supplementary Motor Area | 6 | 4.02 |  |
| -12 | 0 | 74 | Left Supplementary Motor Area | 6 | 3.48 |  |
| -69 | -15 | -21 | Left Middle Temporal Gyrus | 21 | 7.33 | 9564 |
| -68 | -22 | -27 | Left Inferior Temporal Gyrus | 20 | 7.05 |  |
| -70 | -20 | -14 | Left Middle Temporal Gyrus | 21 | 6.63 |  |
| -33 | -78 | -56 | Left Cerebellum Lobule VIIb |  | 4.23 | 495 |
| -40 | -70 | -57 | Left Cerebellum Lobule VIIb |  | 4.03 |  |
| -26 | -72 | -62 | Left Cerebellum Lobule VIII |  | 4.02 |  |
| -4 | -98 | -16 | Left Lingual Gyrus | 18 | 4.61 | 2952 |
| -3 | -100 | -9 | Left Calcarine Fissure | 18 | 4.42 |  |
| -45 | -86 | -15 | Left Inferior Occipital Gyrus | 19 | 4.12 |  |
| 63 | 4 | -22 | Right Middle Temporal Gyrus | 38 | 7.63 | 10669 |
| 66 | -4 | -26 | Right Middle Temporal Gyrus | 21 | 6.71 |  |
| 54 | 18 | -24 | Right Middle Temporal Pole | 38 | 6.68 |  |
| 6 | -66 | -57 | Right Cerebellum Lobule VIII |  | 4.92 | 628 |
| 0 | -58 | -60 | Right Cerebellum Lobule IX |  | 4.35 |  |
| 8 | -58 | -63 | Right Cerebellum Lobule IX |  | 4.29 |  |
| 28 | -75 | -60 | Right Cerebellum Lobule VIII |  | 4.05 | 487 |
| 33 | -80 | -54 | Right Cerebellum Lobule VIIb |  | 3.89 |  |
| 40 | -69 | -58 | Right Cerebellum Lobule VIIb |  | 3.77 |  |
| **Discovery < Replication** | | | | | | |
| 0 | 6 | 4 | Left Caudate | 48 | 3.78 | 457 |
| -10 | -2 | 10 | Left Thalamus | 48 | 3.59 |  |
| -12 | -12 | 20 | Left Thalamus | 48 | 3.55 |  |
| -26 | -28 | 70 | Left Postcentral Gyrus | 4 | 3.89 | 479 |
| -12 | -30 | 78 | Left Paracentral Lobule | 4 | 3.44 |  |
| -20 | -33 | 76 | Left Postcentral Gyrus | 1 | 3.28 |  |
| -68 | -48 | 20 | Left Superior Temporal Gyrus | 39 | 4.28 | 945 |
| -63 | -51 | 39 | Left Supramarginal Gyrus | 39 | 4.23 |  |
| -66 | -50 | 28 | Left Supramarginal Gyrus | 39 | 4.16 |  |
| -30 | -87 | 38 | Left Middle Occipital Gyrus | 19 | 4.63 | 921 |
| -27 | -92 | 32 | Left Superior Occipital Gyrus | 19 | 4.25 |  |
| -27 | -94 | 22 | Left Superior Occipital Gyrus | 19 | 3.9 |  |
| 24 | -10 | 2 | Right Pallidum | 49 | 3.35 | 261 |
| 32 | -14 | 0 | Right Putamen | 49 | 3.16 |  |
| 27 | -12 | 9 | Right Putamen | 49 | 3.01 |  |
| 21 | -30 | 74 | Right Postcentral Gyrus | 4 | 4.81 | 792 |
| 30 | -27 | 72 | Right Precentral Gyrus | 4 | 3.84 |  |
| 36 | -39 | 64 | Right Postcentral Gyrus | 1 | 2.81 |  |
| 21 | -34 | -44 | Right Cerebellum Lobule X | 37 | 4.27 | 263 |
| 15 | -38 | -51 | Right Cerebellum Lobule IX | 37 | 3.56 |  |
| 33 | -88 | 30 | Right Middle Occipital Gyrus | 19 | 4.87 | 3400 |
| 34 | -81 | 44 | Right Superior Occipital Gyrus | 7 | 4.59 |  |
| 22 | -81 | 8 | Right Cuneus | 17 | 4.43 |  |
| **Interaction of Reading Ability by Dataset** | | | | | | |
| **x** | **y** | **z** | **Anatomical Region** | **BA** | **Z** | **kE** |
|  |  |  | None |  |  |  |

*Note: Black font indicates peak coordinates of cluster and its cluster size, and gray font indicates sub-peak coordinates within that cluster. Coordinates indicating anatomical location are in Montreal Neurological Institute (MNI) stereotaxic space. See Figure 2*.
